# Overactive STAT3 drives accumulation of disease-associated CD21^low^ B cells

**DOI:** 10.1101/2021.12.20.473595

**Authors:** Etienne Masle-Farquhar, Timothy Peters, Katherine JL Jackson, Mandeep Singh, Cindy S Ma, Daniel Suan, Gulbu Uzel, Ignatius Chua, Jennifer W Leiding, Kaarina Heiskanen, Kahn Preece, Leena Kainulainen, Michael O’Sullivan, Megan A Cooper, Mikko RJ Seppänen, Satu Mustjoki, Shannon Brothers, Tiphanie P Vogel, Robert Brink, Stuart G Tangye, Joanne H Reed, Christopher C Goodnow

**Author notes:** Senior author (Joanne H Reed, Christopher C Goodnow). These authors contributed equally (Joanne H Reed, Christopher C Goodnow). Lead contact: Christopher C. Goodnow.

## Abstract

Dysregulated STAT3 signalling is correlated with antibody-mediated autoimmunity and B-cell neoplasia, but its effect on B cells is underexplored. Here we address this in children with STAT3 gain-of-function (GOF) syndrome and in mice with STAT3^T716M^, the most common STAT3 GOF syndrome human mutation, or STAT3^K658N^, a dimerization interface mutation responsible for STAT3 GOF syndrome in two children. The main B cell consequence of overactive STAT3 was accumulation of CD19^high^ CD21^low^ atypical memory B cells in humans and of CD21^low^ CD23^low^ B cells in mice resembling age-associated B cells expressing T-bet, CD11c and plasma cell differentiation genes. Overactive STAT3 within B cells increased expression of many genes in the B cell receptor and T cell help pathways, increased the tolerogenic receptor CD22, but opposed B cell tolerance checkpoints and increased formation of T-bet+ B cells upon BCR and CD40 stimulation. These results reveal overactive STAT3 as a central driver of a key class of disease-associated B-lymphocytes in humans and mice.

## INTRODUCTION

Signal Transducer and Activator of Transcription 3 (STAT3) is a cytosolic protein phosphorylated and activated by Janus kinases (JAKs) to regulate gene transcription in response to extracellular hormones and cytokines including interleukin 6 (IL-6) and IL-10 family cytokines, IL- 21, IL-23 and IL-27 (Deenick et al., 2018, O’Shea et al., 2013). Genetic polymorphisms near *STAT3* and genes encoding upstream cytokine receptors increase susceptibility to a range of autoimmune and inflammatory diseases (https://www.ebi.ac.uk/gwas/genes/STAT3). Furthermore, a growing list of inflammatory diseases are now successfully being treated with small molecule JAK inhibitors (Damsky et al., 2021) or monoclonal antibodies that neutralise IL-6 receptor (IL-6R) (Choy et al., 2020) or IL-12/23 signalling. However, therapeutic responses are variable and difficult to predict in individual patients, and rational treatment selection confounded because JAK/STAT3 signalling regulates many cell types and potential targets and the specific consequences of overactive STAT3 remain to be defined.

Notably, several lines of evidence point to B cells as one of the primary cell types affected by dysregulated by JAK/STAT3 signalling. Loss-of-function mutations have shown that STAT3 is cell-autonomously required for B cell germinal centre responses (Ding et al., 2016, Kane et al., 2016), affinity maturation (Kane et al., 2016), plasma cell differentiation (Avery et al., 2010, Deenick et al., 2013) and memory B cell responses (Meyer-Bahlburg et al., 2012, Avery et al., 2010). These observations pose the question of what are the consequences of overactive STAT3 in otherwise normal B cells. STAT3 is constitutively phosphorylated and active in the nucleus in 50% of Activated B Cell-like diffuse large B cell lymphoma (ABC-DLBCL) (Ding et al., 2008) and in >95% of cases of multiple myeloma (Catlett-Falcone et al., 1999), chronic lymphocytic leukemia (CLL; (Hazan-Halevy et al., 2010, Frank et al., 1997)) and mantle cell lymphoma (MCL; (Lai et al., 2003)). In most of these cases it is unclear if active STAT3 is a cause or consequence of the disease, but 6% of DLBCL not otherwise specified (DLBCL-NOS) (Ohgami et al., 2014) and up to 11% of DLBCL (Reddy et al., 2017, Morin et al., 2011, Lohr et al., 2012) have somatic mutations in *STAT3* itself. These mutations cause intrinsic gain-of-function (GOF), often by substituting amino acids in the dimerization interface of the Src Homology 2 (SH2) domain, and thus enhancing dimerization (de Araujo et al., 2019).

Aberrantly high IL-6 serum levels correlate with disease activity in autoimmune thyroid disease (AITD) (Lakatos et al., 1997, Figueroa-Vega et al., 2010), Behçet disease (Akman-Demir et al., 2008), Castleman’s disease (Yoshizaki et al., 1989), rheumatoid arthritis (RA) (Hirano et al., 1988), systemic lupus erythematosus (SLE) (Linker-Israeli et al., 1991) and systemic juvenile idiopathic arthritis (sJIA) (De Benedetti and Martini, 1998). Serum IL-21 levels are elevated in individuals with AITD (Guan et al., 2015), RA (Rasmussen et al., 2010), SLE (Sawalha et al., 2008, Webb et al., 2009) and primary Sjögren’s syndrome (Kang et al., 2011). Anti-IL-6R Tocilizumab is approved (Choy et al., 2020) and clinically beneficial in treating Castleman’s disease (Nishimoto et al., 2000, Nishimoto et al., 2005), RA (Choy et al., 2002), sJIA (Yokota et al., 2005) or the related adult-onset Still’s disease (AOSD) (Kaneko et al., 2018), and JAK inhibitors are approved for the treatment of RA, psoriatic arthritis and JIA, amongst others (Damsky et al., 2021). The variable efficacy of these treatments in some but not all patients with B cell autoimmunity points to the need for increased understanding of the effects of overactive STAT3 in B cells.

Rare children born with heterozygous germline *STAT3* GOF mutations, most frequently a Thr716Met substitution in the C-terminal transactivation (TA) domain, develop early-onset immune dysregulation characterised by lymphadenopathy and splenomegaly, severe autoimmune cytopenias caused by antibodies against blood cell-surface antigens, autoimmune thyroid disease, infectious susceptibility and common variable immune deficiency (CVID) (Flanagan et al., 2014, Haapaniemi et al., 2015, Milner et al., 2015, Bousfiha et al., 2020, Fabre et al., 2019). A crucial open question is how the germline mutations in STAT3 GOF syndrome act: do they dysregulate B cells directly, or secondary to dysregulation of other cell types such as IL-21-producing helper T cells or IL-6- producing macrophages?

A striking cellular abnormality in individuals with CVID and autoimmune cytopenias is the accumulation of atypical CD21^low^ CD23^low^ memory B cells (Isnardi et al., 2010, Warnatz et al., 2002). CD21^low^ CD23^low^ B cell populations variously labelled atypical-memory, IgD/CD27 double-negative or age-associated B cells – referred to collectively here as CD21^low^ CD23^low^ B cells – also accumulate in humans with chronic infections (Benedetto et al., 1992, Moir et al., 2001), severe SLE (Rakhmanov et al., 2009, Wei et al., 2007), RA (Isnardi et al., 2010), primary Sjögren’s syndrome (Saadoun et al., 2013) and in aged mice or mice predisposed to autoimmune disease (Hao et al., 2011, Rubtsov et al., 2011). CD21^low^ CD23^low^ B cells have a gene expression profile intermediate between mature activated B cells and plasmablasts, are enriched for self-reactive B cell receptor (BCR) specificities and poised for antibody production (Charles et al., 2011, Isnardi et al., 2010, Jenks et al., 2018, Rakhmanov et al., 2009, Rubtsov et al., 2011, Russell Knode et al., 2017, Terrier et al., 2011, Scharer et al., 2019).

Here, we show that CD21^low^ CD23^low^ B cells are a major target for dysregulated STAT3 activity. 13/15 individuals with STAT3 GOF syndrome had a striking increase in frequency of circulating CD21^low^ B cells. We find that CD21^low^ CD23^low^ B cells are intrinsically dysregulated to over-accumulate in two different mouse strains: with a K658N mutation in the SH2 domain recurrently mutated in human lymphomas/leukemias, or with the TA domain mutation T716M, the most frequent mutation in STAT3 GOF syndrome (Flanagan et al., 2014, Milner et al., 2015, Fabre et al., 2019). STAT3 GOF dysregulates mouse B cells bearing antibodies against cell surface self-antigens, disrupting immune tolerance to allow their accumulation as CD21^low^ CD23^low^ B cells.

Single-cell mRNA sequencing (scRNA-seq) integrated with chromatin immunoprecipitation sequencing (ChIP-seq) data reveals a landscape of genes dysregulated in B cells by overactive STAT3, including genes recurrently associated with CD21^low^ CD23^low^ B cells like *Tbx21, Zeb2, Fcrl5, Cd22* and *Itgam*, and a suite of genes encoding key elements of the surface immunoglobulin/BCR signalling pathway. Our functional studies demonstrate that STAT3 GOF significantly enhances formation of T-bet^+^ B cells from BCR- and CD40-stimulated follicular B cells. These results reveal overactive STAT3 signalling as a central driver in the formation of CD21^low^ CD23^low^ B-lymphocytes, providing a framework for understanding the pathogenesis and treatment of autoimmunity by modifiers of the STAT3 signalling pathway.

## RESULTS

### Gain-of-function STAT3 drives accumulation of CD21^low^ CD23^low^ B cells

To address the question of whether or not B cells are affected by overactive STAT3, we analysed circulating B cell populations from peripheral blood mononuclear cells (PBMCs) of 15 individuals with STAT3 GOF syndrome, relative to 91 healthy donors. We observed a mean 5.8- fold increase in frequency of circulating B cells with a CD19^high^ CD21^low^ phenotype in STAT3 GOF patients relative to controls. 13 out of 15 patients presented with a significant accumulation of CD19^high^ CD21^low^ B cells, ranging from a 2.9-fold to an 11.9-fold relative increase (**Figure 1A,B**).

**Figure 1.**
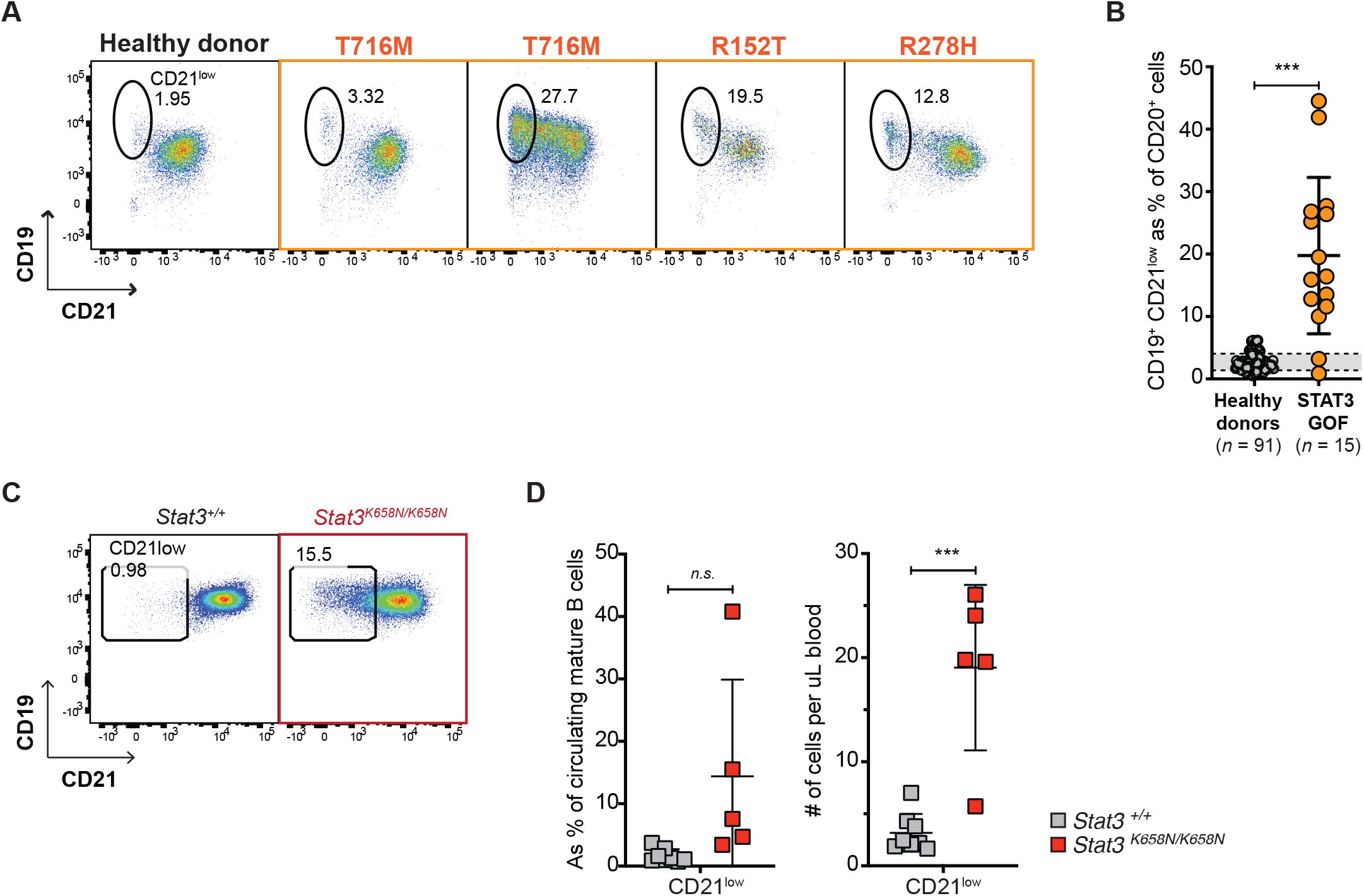
STAT3 GOF mutation increases circulating CD21^low^ B cells in humans and mice. **A,** Representative flow cytometric analysis of CD21 and CD19 expression by circulating CD20+ B cells in peripheral blood mononuclear cells from a healthy donor (left, black outline) or individuals with STAT3 GOF syndrome (orange outline) due to a T716M mutation (*n=2*) or a R152T or R278H mutation. **B,** Percentage of circulating CD20+ B cells with a CD19^high^ CD21^low^ phenotype, in *n*=15 individuals with STAT3 GOF syndrome relative to *n*=91 healthy donors. Comparisons by unpaired *t*-test. * p < 0.05; ** p < 0.01; *** p < 0.001. **C,** Representative flow cytometric analysis of CD21 and CD19 expression by B220^+^ CD95^-^ CD93^-^ mature B cells in the blood of a *Stat3^+/+^* control mouse (left) or a mouse with a homozygous STAT3 gain-of-function K658N mutation (right). **D,** Percentage of CD21^low^ cells in mature B220+ cells and total number per μL blood in individual mice of the indicated genotypes. Comparison by unpaired *t*-test. * p < 0.05; ** p < 0.01; *** p < 0.001. Data are represented as mean ± SD.

To extend this observation, we analysed two strains of mice with *STAT3* GOF mutations generated by CRISPR/*Cas9* genome editing (Masle-Farquhar *et al*., manuscript in preparation – attached to submission for review purposes). The K658N mutation, previously identified in two unrelated children with STAT3 GOF syndrome (Flanagan et al., 2014, Ding et al., 2017), was chosen to model gain-of-function mutations in the STAT3 SH2 domain dimerisation interface. The T716M mutation was chosen to model gain-of-function mutations in the transactivation (TA) domain, and because it is the most frequent germline mutation in STAT3 GOF syndrome (Fabre et al., 2019). *Stat3^K658N/K658N^* mice had a significant increase in frequency of B220^+^ CD93^-^ CD19^+^ CD21^low^ B cells in the blood (**Figure 1C,D**)

*Stat3^+/+^*, *Stat3^K658N/+^* and *Stat3^K658N/K658N^* mice had a comparable frequency of splenic CD19^+^ B cells and relative percentages of CD93^+^ immature and CD93^-^ mature B cells (**Supplementary Figure 1A,B**), as did *Stat3^+/+^*, *Stat3^T716M/+^* and *Stat3^T716M/T716M^* mice (**Supplementary Figure 1C,D**). The most striking B cell abnormality in homozygotes from both strains was an increase in CD21^low^ CD23^low^ mature (CD19^+^ B220^+^ CD93^-^ CD95^-^) B cells. These cells were increased 4-fold as a percentage of mature splenic B cells (**Figure 2A**) and 5.3-fold in total number per spleen (**Figure 2B**), in *Stat3^K658N/K658N^* relative to *Stat3^+/+^* mice. Relative to wild-type controls, *Stat3^T716M/T716M^* mice had a 1.6-fold increase in percentage of CD21^low^ CD23^low^ mature B cells (**Figure 2C**) and 3.2-fold increase in their total number per spleen (**Figure 2D**).

**Figure 2.**
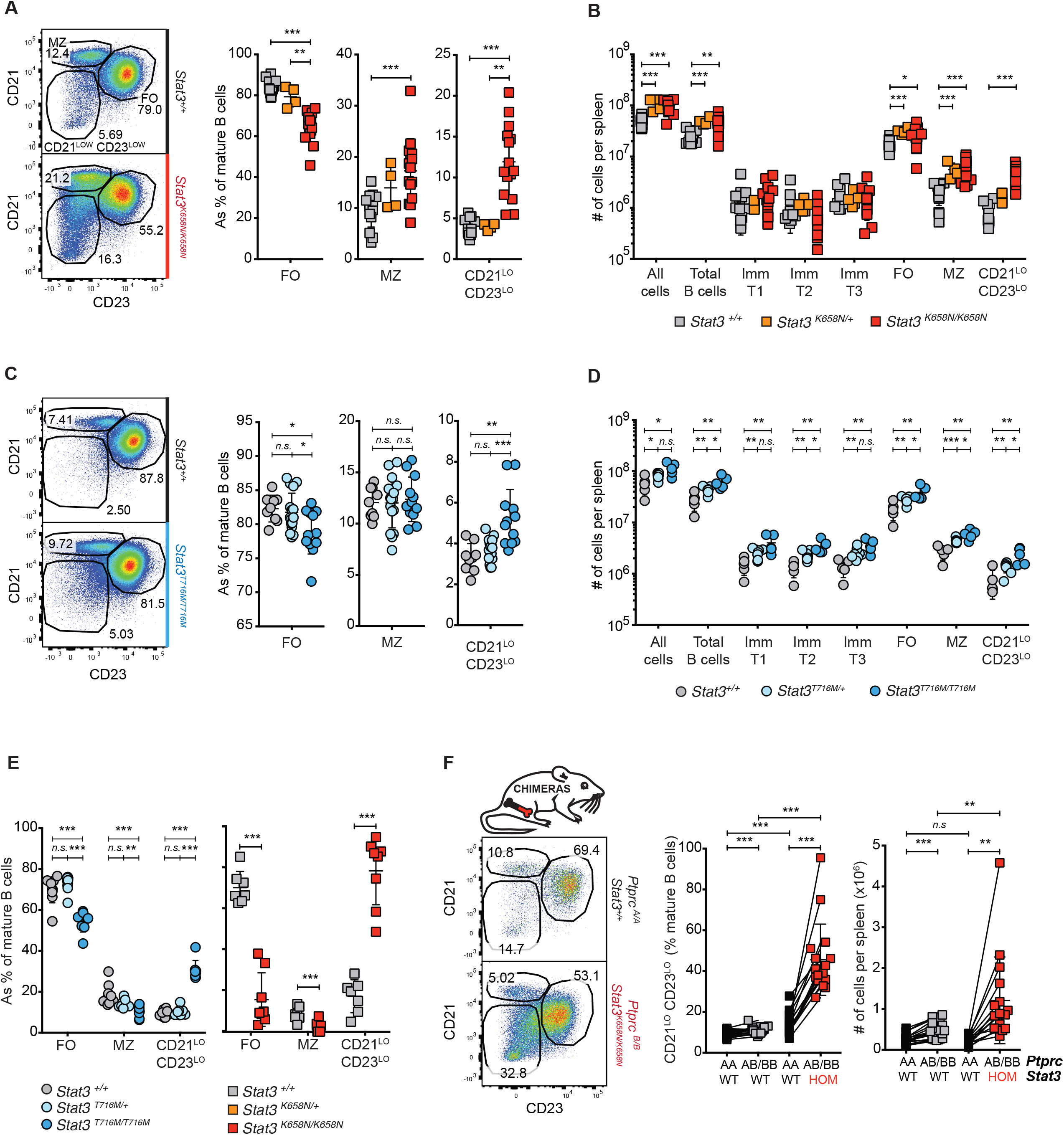
Two different human STAT3 gain-of-function mutations introduced into mice drive accumulation of CD21^low^ CD23^low^ mature B cells. A,B. Mice with STAT3 K658N SH2 domain mutation. A, Representative flow cytometric profiles and percentages of CD19^pos^ CD95^neg^ CD93^neg^ mature splenic B cells gated into the following subsets: FO, follicular CD21^med^ CD23^pos^ ; MZ, marginal zone CD21^high^ CD23^low^ ; and CD21^low^ CD23^low^ B cells. Symbols denote values from individual *Stat3^+/+^* (grey fill), *Stat3^K658N/+^* (orange fill) or *Stat3^K658N/K658N^* (red fill) mice. B, Total number per spleen of CD93^pos^ transitional B cells subsets (T1 CD23^neg^, T2 CD23^pos^ IgM^high^, T3 CD23^pos^ IgM^low^) and CD93^neg^ mature subsets (FO, MZ, CD23^low^ CD21^low^). C,D, Mice with STAT3 T716M transactivating domain mutation. Representative flow cytometric profiles, percentages and total number per spleen of the indicated B cell subsets in *Stat3^+/+^* (grey fill), *Stat3^T716M/+^* (light blue fill) or *Stat3^T716M/T716M^* (dark blue fill) mice. E, Bone marrow chimeras with *Stat3* GOF limited to hematopoietic cells. Percentage of FO, MZ and CD21^low^ CD23^low^ B cells within mature splenic B cells of individual *Stat3^+/+^ Rag1^KO/KO^* mice transplanted with bone marrow of the indicated *Stat3* genotypes. Data representative of *n* > 3 experiments with *n* > 4 mice per group (A-D) or *n* = 3 experiments with *n* > 5 *Rag1^KO/KO^* recipient mice per group (E). Comparisons by *t*-test, corrected for multiple comparisons using the Holm-Sidak method. * p < 0.05; ** p < 0.01; *** p < 0.001. F, Mixed bone marrow chimeras with *Stat3* GOF limited to 50% of hematopoietic stem cells. *Stat3^+/+^ Rag1^KO/KO^* mice were transplanted with bone marrow from *Stat3^+/+^ Ptprc^a/a^* (black fill, WT) donor mice in a 1:1 mixture with bone marrow from a *Stat3^+/+^* (grey fill, WT) or *Stat3^K658N/K658N^* (red fill, HOM) *Ptprc^a/b^* or *Ptprc^b/b^* donor mouse. After staining with PTPRC allele-specific antibodies, representative flow cytometric plots are gated on *Ptprc^a/a^* or *Ptprc^a/b^*/*Ptprc^b/b^* mature B cells, showing percentage of FO, MZ and CD21^low^ CD23^low^ B cells. Symbols in graphs show percentage and number of CD21^low^ CD23^low^ B cells of the indicated genotypes in individual chimeric animals. Solid lines link cells within one chimeric mouse. Data pooled from *n* = 2 experiments with *n* > 4 mice per group. Comparisons within each recipient mouse made by paired t-test. Comparisons between mice made by one-way ANOVA followed by multiple comparison post-tests. * p < 0.05; ** p < 0.01; *** p < 0.001. Data are represented as mean ± SD.

Relative to *Stat3^+/+^* mice, *Stat3^K658N/K658N^* mice though not *Stat3^T716M/T716M^* mice, had an increased percentage of IgM^low^ IgD^high^ mature recirculating B cells in the bone marrow with a CD21^low^ CD23^low^ phenotype (**Supplementary Figure 2A**). *Stat3^K658N/K658N^* mice not only had increased CD21^low^ CD23^low^ B cells in the blood but also in the inguinal lymph nodes (**Supplementary Figure 2B,C**). This was striking, given that CD21^low^ CD23^low^ age-associated B cells are normally excluded from lymph nodes (Cancro, 2020).

Since STAT3 is ubiquitously expressed, we tested whether hematopoietic-restricted GOF *Stat3* mutations were sufficient to drive the abnormal accumulation of CD21^low^ CD23^low^ B cells. *Stat3^+/+^ Rag1^KO/KO^* mice were irradiated and transplanted with bone marrow from *Stat3^T716M/T716M^* or *Stat3^K658N/K658N^* mutant mice or littermate wild-type control donors. *Stat3^+/+^Rag1^KO/KO^* mice transplanted with *Stat3^T716M/T716M^* or *Stat3^K658N/K658N^* bone marrow had a 3.3-fold or 5.fold higher mean percentage of CD21^low^ CD23^low^ mature splenic B cells, relative to control chimeras transplanted with *Stat3^+/+^* marrow (**Figure 2E**).

To test whether *Stat3* GOF acts intrinsically within B cells to drive accumulation of CD21^low^ CD23^low^ cells, or secondary to dysregulation of T cells or other hematopoietic cells, we generated mixed chimeras wherein a fraction of B cells and other hematopoietic cells had mutant *Stat3^K658N/K658N^* and the remainder had normal *Stat3* genes (**Figure 2F**). *Stat3^+/+^ Rag1^KO/KO^* mice were irradiated and transplanted with an equal mixture of “test” bone marrow from *Stat3^K658N/K658N^ Ptprc^a/b^* mice and control bone marrow from *Stat3^+/+^ Ptprc^a/a^* donors. As an additional control, a parallel set of mixed chimeras received *Stat3^+/+^ Ptprc^a/b^* “test” marrow with wild-type *Stat3*. Cells in the reconstituted chimeras were stained with *Ptprc* allele-specific antibodies and an identical gating strategy was applied to *Ptprc^a/b^* and *Ptprc^a/a^* leukocytes. In the same chimeric animals, *Ptprc^a/b^ Stat3^K658N/K658N^* CD21^low^ CD23^low^ B cells accumulated in 5-fold higher mean numbers than *Ptprc^a/a^ Stat3^+/+^* B cells (**Figure 2F**). By contrast, there was no significant difference in the number of CD21^low^ CD23^low^ B cells derived from *Stat3^+/+^ Ptprc^a/b^* marrow, in chimeras where the *Ptprc^a/b^* “test” marrow carried wild-type *Stat3*. Thus overactive STAT3 acts cell-autonomously to drive dysregulated accumulation of CD21^low^ CD23^low^ B cells. Analysis of the mixed chimeras also revealed that *Stat3* GOF acts cell autonomously in splenic follicular and immature CD93^+^ transitional 2 (T2) and transitional 3 (T3) B cells to decrease cell-surface expression of CD21 and CD23 (**Supplementary Figure 2C**).

### Polyclonal accumulation of *Stat3*-mutant CD21^low^ CD23^low^ B cells

Given the above data demonstrating that GOF STAT3 drives cell-intrinsic accumulation of CD21^low^ CD23^low^ B cells, and the recurrence of somatic GOF *STAT3* mutations in B cell lymphomas (Ohgami et al., 2014, Reddy et al., 2017, Morin et al., 2011, Lohr et al., 2012), we analysed clonal diversity by immunoglobulin heavy chain VDJ deep sequencing of mRNA from *Stat3^+/+^* and *Stat3^K658N/K658N^* mature follicular and CD21^low^ CD23^low^ B cells purified by fluorescence-activated cell sorting (FACS). This revealed that the accumulating CD21^low^ CD23^low^ B cells in *Stat3^K658N/K658N^* animals were highly polyclonal and did not represent a clonal neoplasm (**Figure 3A-D**). Compared to follicular B cells, multiple expanded clones were present in the CD21^low^ CD23^low^ B cell population, in both *Stat3* mutant and wild-type mice (**Figure 3A,B**).

**Figure 3.**
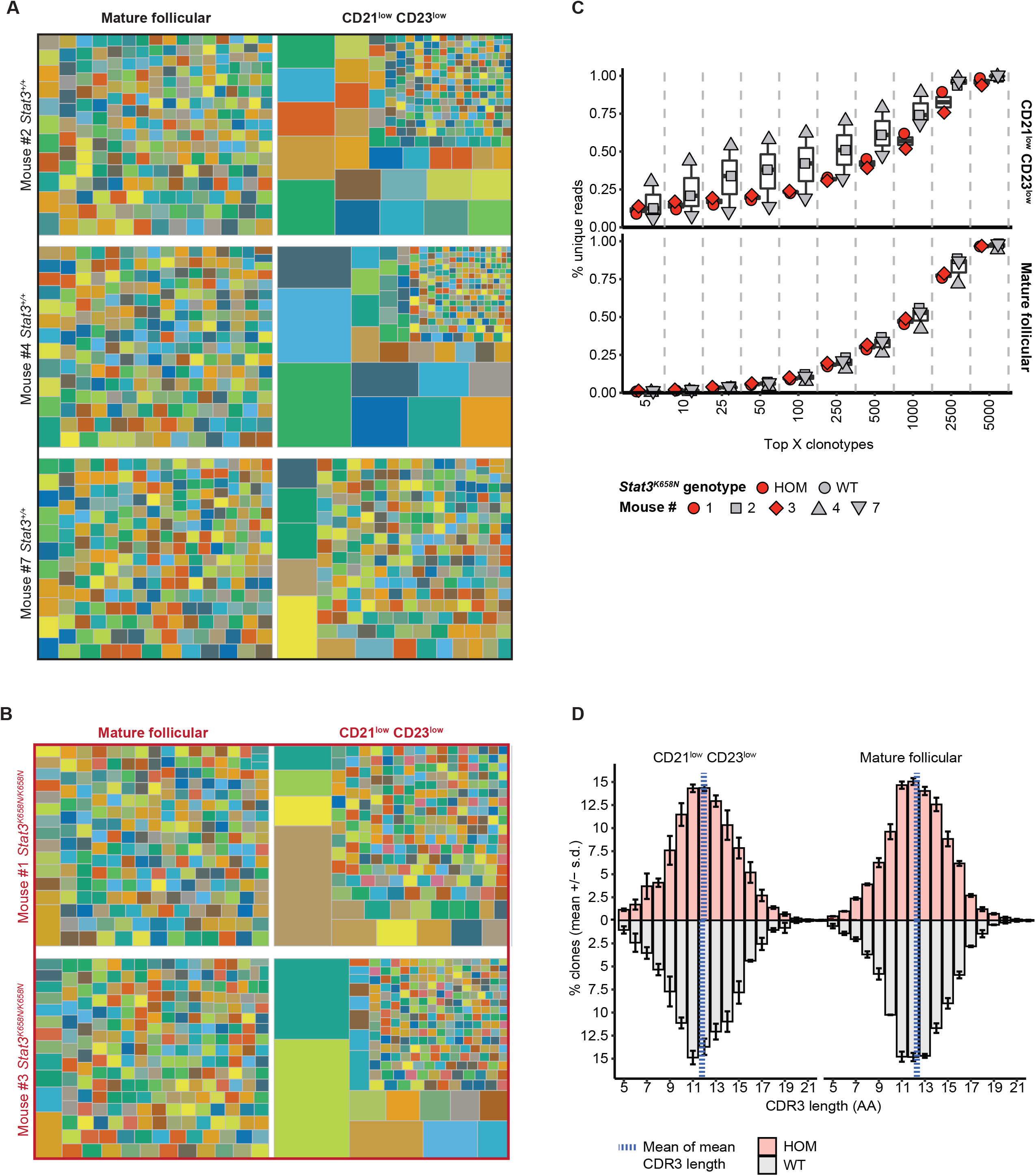
STAT3 mutant CD21^low^ CD23^low^ B cells are clonally diverse. Mature CD93^-^ CD21^low^ CD23^low^ B cells and CD23^+^ follicular B cells were sorted by fluorescence-activated cell sorting (FACS), to prepare independent pools of mRNA from *n*=3 *Stat3^+/+^* and *n*=2 *Stat3^K658N/K658N^* mice. The VDJ region of *Ighm* mRNAs in each pool of cells was amplified by modified 5’ RACE, deep-sequenced, and analysed to identify unique clones based on the CDRH3 nucleotide sequence. **A,B,** Tree maps of the top 250 clones based on number of unique sequence reads in each pool of cells. Each large box represents a single pool of cells of the indicated cell type and genotype. Within each box, the individual-coloured squares correspond to unique clones, with the area proportional to the relative number of unique reads. **C**, Percent of unique reads derived from the top *X* number of clones in CD21^low^ CD23^low^ or follicular B cells from *Stat3^+/+^* (grey fill) or *Stat3^K658N/K658N^* (red fill) mice. **D**, Distribution and mean CDRH3 lengths, in CD21^low^ CD23^low^ or follicular B cells from mice of the indicated genotypes.

CD23^low^ CD21^low^ B cells had comparable CDR3 lengths and hydrophobicity (**Figure 3D**) to mature follicular B cells in *Stat3* wild-type and mutant mice.

### STAT3 gain-of-function mutation allows accumulation of self-reactive B cells *in vivo*

Autoantibody-mediated cytopenias are the most frequent autoimmune manifestation in individuals with STAT3 GOF syndrome (Fabre et al., 2019, Haapaniemi et al., 2015, Milner et al., 2015). This raises the question of how STAT3 GOF affects self-reactive B cells expressing immunoglobulins recognising cell surface autoantigens on blood cells. To address this question, we introduced the *Stat3^T716M^* mutation into a previously established transgenic mouse system (Burnett et al., 2018). In this system, a minor, homogeneous population of SWHEL B cells expressing HyHEL10 immunoglobulin specific for hen egg lysozyme (HEL), can be traced as they develop in chimeric mice expressing a plasma membrane-bound form of HEL with three mutations (mHEL^3X^) as a ubiquitous self-antigen that binds HyHEL10 with micromolar affinity. As illustrated schematically in **Figure 4A**, four sets of chimeric mice were created by bone marrow transplantation to track *Stat3* mutant and wild-type SWHEL B cells developing in the presence or absence of the self-antigen. The recipients comprised *Stat3^+/+^ Ptprc^b/b^* animals of two genotypes: mHEL^3X^-transgenic (Tg) mice and control non-transgenic (non-Tg) mice. Four alternative bone marrow mixtures were transplanted: mHEL^3X^-Tg recipients received polyclonal mHEL^3X^-Tg *Stat3^+/+^ Ptprc^b/b^* marrow mixed with *Ptprc^a/a^ Rag1^KO/KO^* SWHEL bone marrow that was either *Stat3^T716M/T716M^* (HOM) or *Stat3^+/+^* (WT); control non-Tg recipients received polyclonal non-Tg *Stat3^+/+^ Ptprc^b/b^* marrow mixed with *Ptprc^a/a^ Rag1^KO/KO^* SWHEL bone marrow that was either *Stat3^T716M/T716M^* (HOM) or *Stat3^+/+^* (WT).

**Figure 4.**
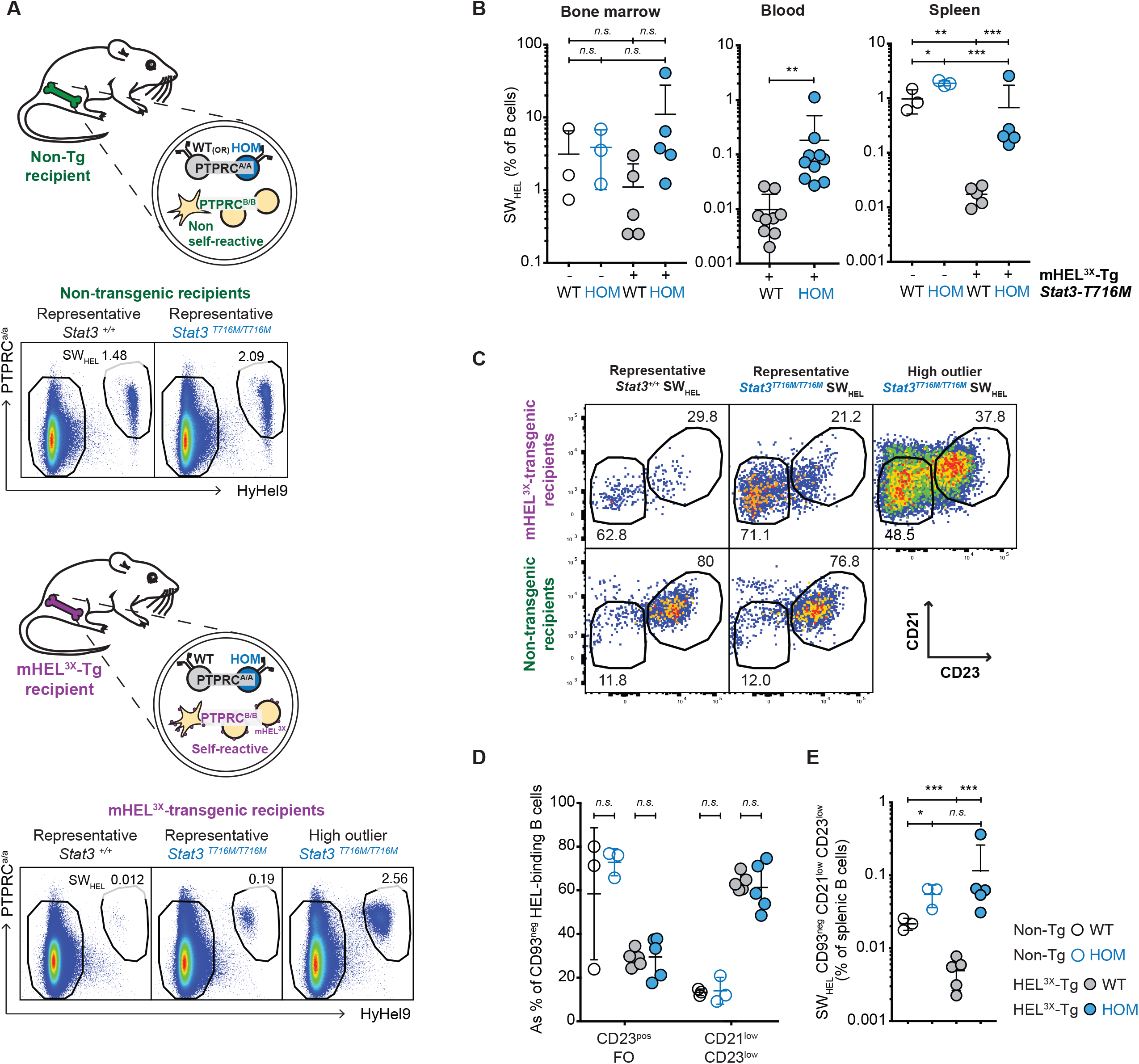
STAT3 gain-of-function increases mature self-reactive CD21^low^ CD23^low^ B cells. **A,** Schematic and representative flow cytometric plots of bone marrow chimeric mice with developing B cells containing a minority of HEL-binding *Ptprc^a/a^* SWHEL B cells that are either *Stat3^+/+^* (WT) or *Stat3^T716M/T716M^* (HOM), and a majority of *Ptprc^b/b^* B cells with diverse immunoglobulins and normal *Stat3* genes. In the lower panels, the marrow transplant recipient and the *Ptprc^b/b^* marrow carry a ubiquitous transgene (Tg) expressing membrane- bound mHEL^3X^ protein, which binds with low affinity to the HyHEL10 immunoglobulin on SWHEL B cells making them self-reactive. In the upper panels, all recipients and donors are non-transgenic (non-Tg), so that the SWHEL B cells are not self-reactive. **B,** Percentage of HEL- binding SWHEL B cells of the indicated *Stat3* genotypes among all B cells in non-Tg or mHEL^3X^- Tg chimeras. **C, D,** Representative plots and percentage of mature CD93^-^ SWHEL B cells in follicular (FO) and CD21^low^ CD23^low^ subsets in individual chimeras of the indicated genotypes. **E**, Percentage of mature CD21^low^ CD23^low^ SWHEL cells among all splenic B cells in individual chimeras. Data are representative of *n* = 2 independent experiments with *n* > 3 recipients per group. After excluding the one high outlier, comparisons made by t-test corrected for multiple comparisons using the Holm-Sidak method. * p < 0.05; ** p < 0.01; *** p < 0.001.

Previous studies have shown that in the presence of mHEL^3X^, self-reactive wild-type SWHEL B cells emigrate from the bone marrow to reach the spleen as immature CD93^+^ surface IgM^low^ T1 and T3 anergic B cells with greatly reduced survival, persisting for only a few days (Burnett et al., 2018). Consistent with this, *Stat3^+/+^* SWHEL B cells were present in comparable frequencies in the bone marrow but were much less frequent in the spleen of mHEL^3X^-Tg chimeras compared to non-Tg controls where no self-antigen was present (**Figure 4A,B**). While *Stat3^T716M/T716M^* SWHEL B cells were also present in comparable frequencies to their wild-type SWHEL counterparts in the bone marrow, these cells were 10-fold more frequent in the spleen and blood of mHEL^3X^-Tg recipients compared to *Stat3^+/+^* self-reactive SWHEL B cells (**Figure 4A,B**). Irrespective of their *Stat3* genotype, 60% of mature CD93-negative SWHEL B cells in mHEL^3X^-Tg chimeras had a CD23^low^ CD21^low^ phenotype, as opposed to approximately 10% of mature SWHEL B cells in non-Tg chimeras where the B cells were not self-reactive (**Figure 4C,D**). The *Stat3^T716M^* mutation, limited only to the SWHEL B cells in these chimeric animals, increased the median frequency of self-reactive CD21^low^ CD23^low^ mature B cells 11-fold, as a percentage of all B cells in the spleen (**Fig 4E**). Thus STAT3 GOF acts cell autonomously to oppose B cell tolerance checkpoints that normally prevent nascent B cells recognizing blood cell surface autoantigens from accumulating as mature B cells with an atypical memory B cell phenotype.

### Overactive STAT3 alters expression of STAT3-binding genes in B cells

To define the transcriptional state of this dysregulated mature B cell population, we performed single-cell RNA sequencing analysis of mature follicular and CD21^low^ CD23^low^ B cells sorted from *Stat3^+/+^* and *Stat3^K658N/K658N^* mice (**Figure 5A**). Cell type and *Stat3* genotype were the principal components of gene expression variation between the sorted cell populations (**Figure 5B**).

**Figure 5.**
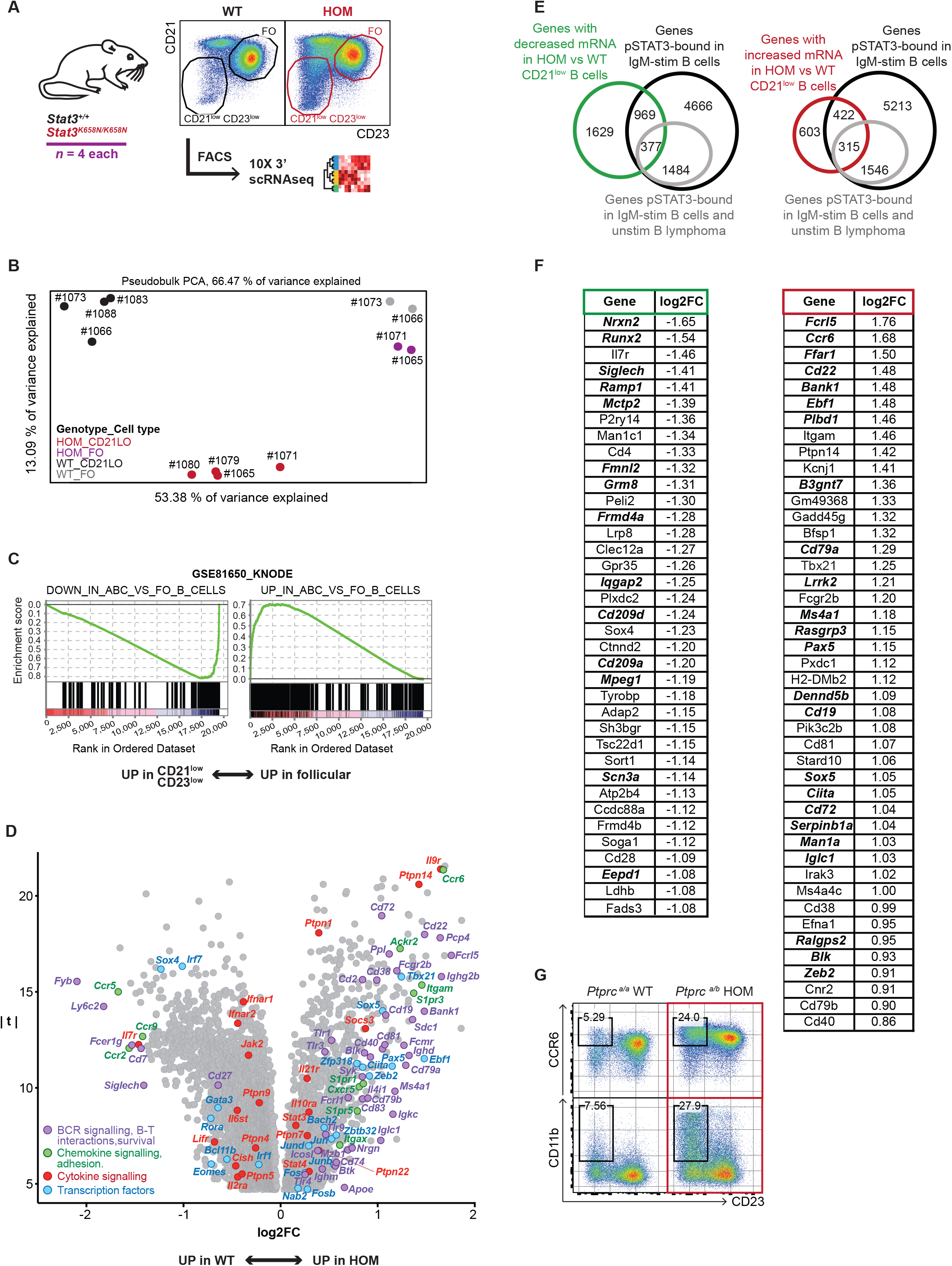
STAT3 gain-of-function changes gene expression in CD21^low^ CD23^low^ B cells. **A**, Workflow of fluorescence-activated cell sorting and single-cell RNA sequencing of mature CD93^-^ B220^+^ CD95^-^ spleen B cells of the CD21^low^ CD23^low^ (*n*=4 donors from each genotype) and follicular (*n*=2 donors from each genotype) subsets of *Stat3^K658N/K658N^* (HOM) and *Stat3^+/+^* (WT) mice. **B**, Pseudobulk principle component analysis of the indicated B cell populations, sorted from individual mice denoted by numbers. **C,** Comparison of sorted CD21^low^ CD23^low^ and FO B cells by gene set enrichment analysis, showing results for GSE81650 comprising a set of mRNAs previously shown (Russell Knode et al., 2017) to be decreased (left panel) or increased (right panel) in age-associated B cells (ABC, CD21^-^ CD23^-^ B220^+^ CD93^-^ CD43^-^ cells) compared to FO B cells (CD21^+^ CD23^+^ B220^+^ CD93^-^ CD43^-^). **D,** Comparison of HOM vs WT CD23^low^ CD21^low^ B cells. Volcano plot of differentially expressed genes with log2FC≠0 and FWER<0.05. Exemplar genes are labelled. **E, F,** Venn diagrams and list of most differentially expressed genes at intersection between genes binding phosphorylated STAT3 (pSTAT3) by chromatin immunoprecipitation sequencing (ChIP-seq) of BCR-stimulated primary human peripheral naïve B cells (Lu et al., 2019) (black circles), or also pSTAT3 bound in unstimulated TMD8 B lymphoma cells (grey circles) (Lu et al., 2019) and genes with increased (red circle) or decreased (green circle) expression in HOM relative to WT CD21^low^ CD23^low^ B cells. FWER of intersection by Fisher exact test: increased genes, p = 4.81 x 10^-54^ (odds ratio 3.30); decreased genes, p = 6.60 x 10^-8^ (odds ratio 1.47). Genes that were also pSTAT3-bound by ChIP-seq of unstimulated TMD8 lymphoma cells (Lu et al., 2019) are in bold and italicised.

Compared to follicular B cells of the same *Stat3* genotype, CD21^low^ CD23^low^ B cells differentially expressed 9680 and 6820 genes in *Stat3^+/+^* or *Stat3^K658N/K658N^* mice, respectively (**Supplementary Table 1**). Gene set enrichment analysis (GSEA) comparing the follicular and CD21^low^ CD23^low^ B cell populations, regardless of their *Stat3* genotype, revealed highly skewed expression of most of the genes in sets previously defined as differentially expressed by age-associated B cells in mice (B220^+^ CD93^-^ CD43^-^ CD21^-^ CD23^-^ in GSE81650 (Russell Knode et al., 2017); B220^+^ CD19^+^ CD11b^+^ in GSE28887 (Rubtsov et al., 2011)) (**Figure 5C**, **Supplementary Figure 3A, Supplementary Table 2**). This included genes distinguishing CD21^low^ CD23^low^ B cells in multiple studies in humans and mice (Charles et al., 2011, Jenks et al., 2018, Lau et al., 2017, Rakhmanov et al., 2009, Rubtsov et al., 2011, Russell Knode et al., 2017, Scharer et al., 2019): increased expression of *Gas7, Ccr2, Fcer1g, Fyb, Runx2, Ahnak, Lair1, Lgals1, Itgax* (CD11c)*, Cxcr3, Zeb2, Zbtb20, Ahnak2, Zbtb32, Itgam* (CD11b)*, Prdm1* (BLIMP-1) and *Tbx21* (T-bet); decreased expression of *Fcer2a* (CD23)*, Cr2* (CD21)*, Crisp3, Icosl, Fcrl1, CD40, Bach2, Ptaj, Il4ra* (**Supplementary Figure 3B, Supplementary Table 1**). Increased expression of *Prdm1, Zbtb20, Zbtb32, Mzb1* and *Zeb2*, and decreased *Bach2* are consistent with CD21^low^ CD23^low^ B cells being primed for plasma cell differentiation (Lau et al., 2017).

To identify genes with dysregulated expression in *Stat3*-mutant CD21^low^ CD23^low^ B cells compared to their wild-type counterparts, we compared single-cell RNA expression by these cells sorted from *Stat3^K658N/K658N^* (HOM) relative to *Stat3^+/+^* (WT) mice (*n* = 4 mice per group) (**Figure 5A,D**). *Stat3^K658N/K658N^* CD21^low^ CD23^low^ B cells differentially expressed 4315 genes relative to *Stat3^+/+^* CD21^low^ CD23^low^ B cells (**Supplementary Table 3**).

1340 genes were increased in *Stat3^K658N/K658N^* CD21^low^ CD23^low^ B cells, many encoding mRNAs and proteins previously found to be increased in disease-associated CD21^low^ CD23^low^ B cells (*Fcrl5, Cd22, Itgam, Sdc1, Cd79a, Ms4a1, Cd72, Cd40, Ciita, Blk, Ilri1, Cd79b, Syk, Itgax, Btk*), involved in cytokine signalling (*Il9r, Ptpn14, Socs3, Ptpn1, Ptpn22, Il10ra, Il21r, Ptpn7, Stat4, Stat3*), chemokine receptors/adhesion molecules (*Ccr6, S1pr3, Ackr2, S1pr1, Cxcr5, S1pr5*), and transcription factors (*Ebf1, Tbx21, Sox5, Pax5, Zeb2, Zfp318, Zbtb32, Junb, Jun, Bach2, Jund, Fosb, Fos, Nab2*) (**Figure 5D**).

2975 genes were decreased in *Stat3^K658N/K658N^* relative to *Stat3^+/+^* CD21^low^ CD23^low^ cells, many encoding molecules involved in cytokine signalling (*Il7r, Lifr, Ifnar2, Cish, Il6st, Ifnar2, Il2ra, Ifnar1, Ptpn5, Jak2, Ptpn9, Ptpn4*), chemokine receptors/adhesion (*Ccr5, Ccr2, Ccr9*), and transcription factors (*Sox4, Irf7, Eomes, Rora, Gata3, Bcl11b, Irf1*) (**Figure 5D**).

To analyse which of the differentially expressed genes in *Stat3*-mutant CD21^low^ CD23^low^ B cells correspond to genes bound by STAT3 in B cells, we used recently published datasets of chromatin immunoprecipitation sequencing (ChIP-seq) of phosphorylated STAT3 (pSTAT3)-bound genes in primary human naïve peripheral blood B cells stimulated with anti-IgM (Lu et al., 2019) (**Supplementary Table 4**). Given that STAT3 is constitutively phosphorylated in many human B cell lymphomas, as a stringent test we analysed which of the pSTAT3 targets in IgM-stimulated B cells also had pSTAT3 bound in unstimulated TMD8 diffuse large B cell lymphoma cells (Lu et al., 2019). Among the 1340 and 2975 increased and decreased genes in *Stat3^K658N/K658N^* relative to *Stat3^+/+^* CD21^low^ CD23^low^ B cells, 737 (55%) and 1346 (45%), respectively, corresponded to pSTAT3 targets in IgM-stimulated human peripheral naïve B cells (**Figure 5E**, **Supplementary Table 4**) and 315 (24%) and 377 (12.7%), respectively, were also pSTAT3 targets in TMD8 lymphoma cells (**Supplementary Figure 3C, Supplementary Table 4**). Among the pSTAT3 target genes in human naïve B cells were 44 and 37 of the 100 genes with the greatest increase or decrease, respectively, in log2 expression fold-change in *Stat3^K658N/K658N^* relative to *Stat3^+/+^* CD21^low^ CD23^low^ B cells (**Figure 5F**), including: *Ccr6* and *Itgam* (CD11b, Complement receptor 3, Mac-1), which are upregulated on memory B cells and promote their migration to sites of inflammation (Elgueta et al., 2015, Kawai et al., 2005, McHeyzer-Williams et al., 2000); genes whose expression recurrently defines CD21^low^ CD23^low^ age-associated/atypical memory B cells including *Itgam, Tbx21* (T-bet) and *Fcrl5*; and genes governing BCR signalling including *Cd22, Cd79a, Ms4a1* (CD20)*, Cd19,* and *Cd72*.

We extended the single cell mRNA sequencing analysis by measuring CCR6 and CD11b protein expression on the surface of B cell populations from mixed chimeras generated as above. Flow cytometry confirmed dramatically increased mean cell-surface CCR6 on *Stat3^K658N/K658N^* and *Stat3^T716M/T716M^* CD21^low^ CD23^low^ B cells relative to their *Stat3^+/+^* counterparts, and also on splenic immature T3, mature follicular, marginal zone B cell subsets (**Supplementary Figure 4A, B**). Thus overactive STAT3 acts directly within B cells to dramatically increase CCR6, a critical G-protein coupled receptor responding to chemokine CCL20 produced by inflamed endothelium and epithelium that is necessary for memory B cell recall responses and localisation to sites of inflammation (Elgueta et al., 2015, Schutyser et al., 2003). The same was true for CD11b and to a lesser extent CD11c on the surface of *Stat3^K658N/K658N^* relative to *Stat3^+/+^* B cell subsets (**Supplementary Figure 4C,D**). CCR6 was homogeneously increased on *Stat3*-mutant CD21^low^ CD23^low^ B cells, whereas only a fraction of these cells expressed high CD11b (**Figure 5G**, **Supplementary Figure 4**). CD11b^+^ CD23^low^ CD21^low^ B cells accumulated in the blood and spleen of *Stat3^K658N/K658N^* mice, and to a lesser extent in *Stat3^T716M/T716M^* mice (**Supplementary Figure 4E, F**), and many of the accumulating CD11b^hi^ cells also expressed CD11c (**Supplementary Figure 4E**).

### *Stat3*-mutant CD21^low^ CD23^low^ B cells have aberrantly high expression of BCR signalling molecules

Gene set enrichment analysis (GSEA) of mRNAs differentially expressed in *Stat3^K658N/K658N^* relative to *Stat3^+/+^* CD21^low^ CD23^low^ B cells revealed the most significant GO term was “B cell receptor signalling pathway” (FDR q=0.001), and the “KEGG B cell receptor signalling pathway” was one of the most significant curated terms (FDR q=0.012) (**Figure 6A**, **Supplementary Table 5**). Of the 25 leading edge genes in the KEGG BCR signalling pathway that were significantly increased in *Stat3^K658N/K658N^* relative to *Stat3^+/+^* CD21^low^ B cells, all 25 corresponded to ChIPseq pSTAT3-bound genes in IgM-stimulated human B cells, and 15 were also pSTAT3 bound in TMD8 lymphoma cells (Lu et al., 2019) (**Figure 6B**, **Supplementary Table 5**). *Cd22*, *Ighd* and *Cd79a* had the highest fold change among these gene-sets, and were validated by flow cytometry showing cell-autonomously increased cell surface CD22 and IgD in *Stat3^K658N/K658N^* and *Stat3^T716M/T716M^* CD21^low^ B cells (**Figure 6C**, **Supplementary Figure 5A**).

**Figure 6.**
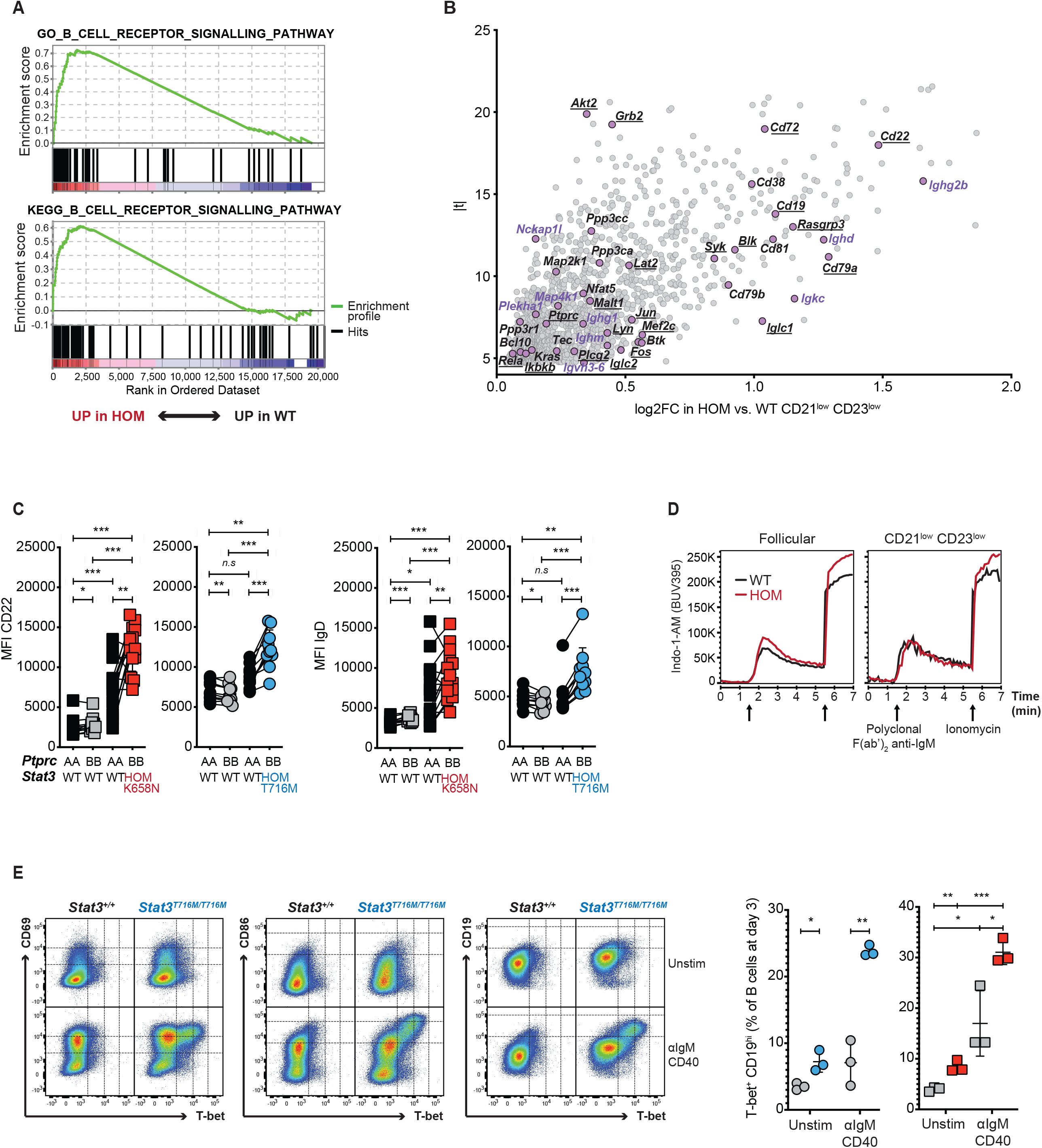
STAT3 gain-of-function dysregulates the response to BCR and CD40 stimulation. **A,** Gene set enrichment analysis (GSEA) showing results from single cell RNA sequencing analysis comparing *Stat3^K658N/K658N^* (HOM) to *Stat3^+/+^* (WT) CD21^low^ CD23^low^ B cells for the 50 genes in the Gene Ontology (GO) B cell receptor signalling pathway (top, FDR q=0.001, FWER p = 0.001) and 67 mostly non-overlapping genes in the KEGG B cell receptor signalling pathway (bottom, FDR q=0.011, FWER p = 0.15). Plots show rank-ordered genes (x axis) and enrichment scores for genes in each set (y axis). Details are in **Supplementary Table 5**. **B**, Volcano plot of mRNA expression log2 fold-change (log2FC) versus moderated *t*-statistic for increased genes (log2FC>0 and FWER<0.05) in HOM relative to WT CD21^low^ CD23^low^ B cells. Purple circles denote genes in the top 25 leading edge of the GO and KEGG B cell receptor signalling pathway GSEAs. Black text denotes genes pSTAT3-bound in anti-IgM stimulated human naïve B cells (Lu et al., 2019). Underlined black text denotes genes also pSTAT3-bound in TMD8 lymphoma cells (Lu et al., 2019). Purple text denotes genes pSTAT3-bound in neither population. **C,** Mean fluorescence intensity (MFI) of CD22 and IgD staining on *Stat3* mutant and wild-type CD21^low^ CD23^low^ B cells in mixed chimeras. **D**, Mean fluorescence intensity of calcium-bound Indo-1-AM at 400 nm in splenic follicular and CD21^low^ CD23^low^ B cells from *Stat3^K658N/K658N^* (HOM) and *Stat3^+/+^* (WT) mice, preceding and following addition of polyclonal F(ab’)2 anti-mouse IgM (first arrow) and ionomycin (second arrow). **E,** CD23^+^ follicular B cells, isolated by magnetic activated cell-sorting from the spleen of individual mice of the indicated genotypes, were cultured for 3 days unstimulated or stimulated with anti-IgM F(ab)’2 and CD40, and analysed by flow cytometry. Right panels show % of T-bet^+^ CD19^hi^ B cells in individual cultures. Each symbol is the average of a technical duplicate, for follicular B cells obtained from an individual *Stat3^+/+^* (grey fill), *Stat3^T716M/T716M^* (blue fill) or *Stat3^K658N/K658N^* (red fill) mouse. Comparisons made by t-test, corrected for multiple comparisons using the Holm-Sidak method. * p < 0.05; ** p < 0.01; *** p < 0.001. Data are represented as mean ± SD.

The impact of these gene expression changes was analysed by stimulating the BCR with F(ab’)2 anti-mouse IgM *in vitro* (**Figure 6D**). Despite lower levels of cell-surface IgM expression (**Supplementary Figure 5B,C**), CD21^low^ CD23^low^ B cells from *Stat3^K658N/K658N^* mice had no discernable difference in their intracellular calcium response to IgM stimulation compared to wild-type control cells, and mutant follicular B cells displayed a subtle increase (**Figure 6D**). Given the evidence that activated follicular B cells become CD21^low^ CD23^low^ B cells, we enriched for splenic CD23^+^ follicular B cells from *Stat3^K658N/K658N^*, *Stat3^T716M/T716M^* or *Stat3^+/+^* mice and cultured them in complete RPMI (cRPMI) with F(ab’)2 anti-mouse IgM and a stimulatory anti-CD40 monoclonal antibody, or cRPMI alone (unstim), for 3 days (**Figure 6E**). In response to IgM and CD40 stimulation, both STAT3 GOF mutations dramatically increased the formation of cells expressing intracellular T-bet and high levels of cell-surface CD86 and CD19.

## DISCUSSION

The findings presented here address key questions about the role of dysregulated STAT3 signalling in B cell lymphomas and in antibody-mediated autoimmune disease. Constitutively phosphorylated STAT3 and *STAT3* GOF mutations are frequent in human B cell leukemias and lymphomas, and the findings here demonstrate that overactive STAT3 in B cells is sufficient to drive aberrant accumulation of CD21^low^ B cells in mice and humans. In mice, STAT3 GOF drives polyclonal expansion of atypical CD21^low^ CD23^low^ B cells with dramatically dysregulated gene expression but is insufficient to cause formation of clonal B cell tumours, which presumably require additional driver mutations. Our results reveal that overactive STAT3 acts intrinsically within B cells to dysregulate expression of key homing receptors, genes in the T cell help and BCR signalling pathways. With respect to autoimmune disease, STAT3 GOF opposes B cell tolerance checkpoints, exaggerates the response to B cell receptor stimulation and drives polyclonal accumulation of CD21^low^ CD23^low^ mature B cells with a gene expression profile closely resembling age-associated B cells and atypical memory B cells/double negative memory B cells found in autoimmune syndromes in mice and humans.

The results here are directly relevant to understanding the autoimmune anemia, thrombocytopenia and other cytopenias that occur in a high proportion of individuals with germline *STAT3* GOF mutations. The T716M mutation studied here is the most common mutation causing childhood STAT3 GOF syndrome (Fabre et al., 2019), while the K658N mutation has been reported to cause STAT3 GOF syndrome in several unrelated individuals (Flanagan et al., 2014, Ding et al., 2017). Despite carrying STAT3 mutations in every cell in their body from birth, affected individuals only develop autoimmune cytopenias after a delay varying from months to years. We show here that most STAT3 GOF syndrome patients have increased frequencies of circulating CD21^low^ B cells. By tracing SWHEL B cells bearing BCRs with low affinity for a defined self-antigen displayed on the surface of erythrocytes and leukocytes, the results here show that STAT3^T716M^ acts within self-reactive B cells to interfere with the checkpoint that normally prevents their accumulation as mature follicular B cells and CD21^low^ CD23^low^ B cells.

The single cell RNA sequencing performed here reveals the circuitry of STAT3-regulated genes within CD21^low^ CD23^low^ B cells. These cells accumulate in the majority of individuals that we tested (with a range of *STAT3* GOF mutations) and in people with systemic lupus (Wehr et al., 2004), rheumatoid arthritis (Isnardi et al., 2010), Sjogren’s syndrome (Saadoun et al., 2013), cryoglobulinemic vasculitis (Charles et al., 2011), and autoimmune cytopenias associated with common variable immunodeficiency (Warnatz et al., 2002). CD21^low^ CD23^low^ B cells in *Stat3^K658N/K658N^* mice had increased expression of 1340 genes compared to their cellular counterparts with wildtype *Stat3*, more than half corresponding to genes with bound phosphorylated STAT3 in IgM-stimulated human B cells (Lu et al., 2019). At the top of this list are widely studied markers of CD21^low^ B cells: T-bet *(Tbx21*), CD11b (*Itgam*), CD22 and FCRL5, and a marker of normal and CD21^low^ memory B cells, CCR6.

T-bet is expressed by CD11b^+^ CD11c^+^ age-associated CD21^low^ B cells (Rubtsov et al., 2011) and is partially required for accumulation of these cells in SLE1,2,3 autoimmune mice (Rubtsova et al., 2017) but not in *Was*-mutant autoimmune mice (Du et al., 2019). In interferon- and TLR9/7- stimulated mouse B cells, T-bet activates sterile transcripts of the *Ighg2a/b/c* locus to promote T cell-independent switching to produce pathological IgG2a/b/c antibodies (Peng et al., 2002). Thus dysregulated induction of *Tbx21* may account for our finding that *Ighg2b* was one of the most increased mRNAs in *Stat3* mutant CD21^low^ B cells. Enforced T-bet expression induces CD11b and CD11c expression by B cells *in vitro* (Rubtsova et al., 2013). Here, we showed that two different *Stat3* GOF mutations significantly increase accumulation of T-bet^+^ B cells in response to IgM and CD40 stimulation *in vitro*. These results support the conclusion that overactive STAT3 acts in part by inducing T-bet in IgM-stimulated B cells, which in turn contributes to driving and shaping the accumulation of CD21^low^ CD23^low^ B cells with a transcriptional status poised for plasma cell differentiation.

*Cd22* was the most increased gene among the KEGG BCR signalling gene set in CD21^low^ B cells with STAT3 GOF. CD22 inhibits IgM signalling and modulates surface IgM levels upon CD22 phosphorylation by LYN kinase and recruitment of SHP-1 (*Ptpn6*) tyrosine phosphatase, and heterozygous loss-of-function mutations establish that *Cd22* and *Lyn* are rate-limiting in this negative feedback (Cornall et al., 1998, Doody et al., 1995). Increased expression of *Cd22* and *Lyn* by STAT3^GOF^ mutant CD21^low^ B cells are therefore predicted to dampen IgM-induced calcium responses, in contrast to the increased expression of *Cd79a*, *Syk*, *Btk* and *Plcg2* mRNAs encoding signalling molecules that promote the IgM-induced calcium response. Despite significantly lower levels of cell-surface IgM expression by *Stat3^GOF^* CD21^low^ CD23^low^ B cells, these cells had comparable intracellular calcium responses to anti-IgM stimulation, indicating that per-IgM molecule, *Stat3*-mutant CD21^low^ CD23^low^ B cells may be hyper-responsive to IgM stimulation.

Since CD22 binding to sialic acid on self-proteins and cells promotes tolerance, and deficiency of *Cd22*, *Lyn* or *Ptpn6* promotes autoimmunity (Meyer et al., 2018), the finding here that *Cd22* and *Lyn* are increased by overactive STAT3 and correspond to phospho-STAT3 bound genes in IgM-stimulated B cells and DLBCL illuminates a checkpoint circuit to counter B cell stimulation by antigens and STAT3-activating cytokines. Speculatively, counter-regulation by CD22 recognizing sialic acids on other blood cells may contribute to why it often takes several years before autoimmune cytopenias develop in children born with STAT3 gain-of-function mutations.

Regardless of CD22 counter-regulation, BCR stimulation coupled with CD40 stimulation nevertheless elicited dramatically increased accumulation of T-bet+ B cells when these cells carried either of the homozygous STAT3 GOF mutations. In mice, accumulation of T-bet+ CD21^low^ CD23^low^ age-associated B cells requires cognate T cell help, based on their failure to develop in CD40 ligand-deficient mice and from adoptive transfer experiments using MHC-II- or CD40- deficient follicular B cells (Russell Knode et al., 2017). Similarly, individuals with germline missense mutations in *CD40* or *CD40LG* have significantly reduced frequencies of circulating CD21^low^ B cells with age (Keller et al., 2021). It is notable that pSTAT3 ChIP-seq target genes with increased mRNA in *Stat3^GOF^* mutant CD21^low^ CD23^low^ B cells included *Cd40* itself, *Ciita* encoding the transcription factor promoting MHC class II expression, *Cd74* encoding the MHC-II invariant chain, *Icosl* encoding the ligand for ICOS costimulation of T follicular helper cells, and *Il21r* encoding the receptor for a key B cell activating cytokine made by T follicular helper cells.

Collectively, the results here reveal that dysregulated STAT3 amplifies the formation of CD23^low^ CD21^low^ B cells in mice and humans, acting at many points in B cell activation (**Figure 7**). This knowledge provides a detailed framework for solving the variability and adverse events in treating autoimmune diseases or B cell neoplasia with small molecule or monoclonal antibody inhibitors of the cytokine-JAK-STAT3 pathway.

**Figure 7.**
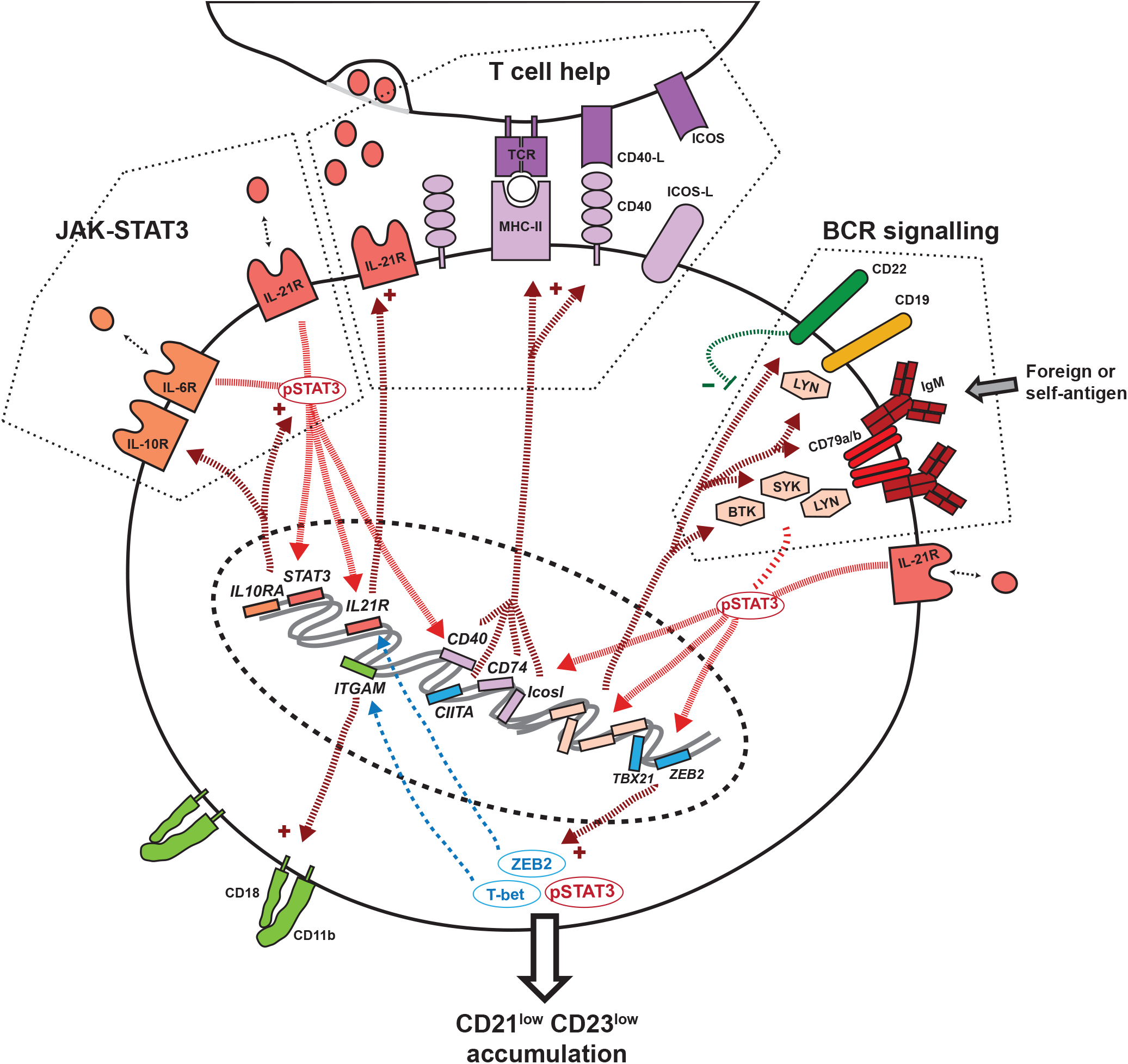
Schematic diagram of pathways dysregulated by overactive STAT3 in CD21^low^ CD23^low^ B cells. Each gene in this schematic diagram belongs to the intersection of: genes pSTAT3-bound following ChIP-seq in human IgM-stimulated peripheral naïve B cells (Lu et al., 2019), and genes with increased mRNA levels in *Stat3^K658N/K658N^* relative to *Stat3^+/+^* B cells (Supplementary Table 4).

## ACKNOWLEDGEMENTS

This work was supported by National Health and Medical Research Council (NHMRC) Program (1113904, to C.C.G.) and Fellowship (1081858, to C.C.G.) grants and by The Bill and Patricia Ritchie Foundation. We thank the Garvan-Weizmann Centre for Cellular Genomics (Garvan Institute of Medical Research, Sydney, Australia) and the Garvan Biological Testing Facility, for providing technical services. We thank Danielle T. Avery, Kathryn J. Payne and Geetha Rao for performing flow cytometry of human samples.

## AUTHOR CONTRIBUTIONS

E.M-F designed and performed experiments; T.J.P and K.J.J designed and performed bioinformatic and statistical analyses of single cell mRNA and VDJ deep sequencing; R.B. and C.C.G designed and generated CRISPR/Cas9 *Stat3*-mutant mouse models; E.M-F and M.S generated BCR deep sequencing libraries; C.S.M, D.S, G.U, I.C, J.W.L, K.H, K.P, L.K, M.O, M.A.C, M.R.J.S, S.M, S.B, S.G.T obtained patient peripheral blood samples; E.M-F, J.H.R and C.C.G interpreted experiments and wrote the manuscript.

## STAR METHODS

### RESOURCE AVAILABILITY

#### Lead contact

Further information and requests for resources and reagents should be directed to and will be fulfilled by the Lead Contact, Christopher C. Goodnow.

#### Materials availability

This study did not generate new unique reagents.

#### Data and code availability

- Single-cell RNA-seq data have been deposited at ENA and are publicly available as of the date of publication. The accession number for these data is European Nucleotide Archive (ENA): PRJEB49382, as listed in the Key Resources table.
- Bioinformatic workflows used to analyse the data in this study are available upon request.
- Any additional information required to reanalyze the data reported in this paper is available from the lead contact upon request.

### EXPERIMENTAL MODEL AND SUBJECT DETAILS

#### Human subjects

This study was approved by the respective ethics review boards of the participating institutes, including the Ethics committees of Sydney Local Health District RPAH Zone Human Research Ethics Committee and Research Governance Office, Royal Prince Alfred Hospital, Camperdown, NSW, Australia (Protocol X16-0210/LNR/16/RPAH/257); the South East Sydney Local Health District Human Research Ethics Committee, Prince of Wales/Sydney Children’s Hospital, Randwick, NSW, Australia (Protocol HREC/11/POWH/152). Written informed consent for genetic investigations and immunological analyses, as well as publication of data, was obtained from each family.

#### Mouse handling, housing and husbandry

All mouse handling and experimental methods were performed in accordance with approved protocols of the Garvan Institute of Medical Research/St Vincent’s Hospital Animal Ethics Committee. All mice were bred and maintained in specific pathogen-free conditions, and housed with littermates, at Australian BioResources (ABR; Moss Vale, Australia) or at the Garvan Institute of Medical Research Biological Testing Facility (BTF). 7 to 15 week-old wild-type and mutant mice, or wild-type and mutant bone marrow recipients, were sex- and age-matched. All experiments conformed to the current guidelines from the Australian Code of Practice for the Care and Use of Animals for Scientific Purposes. Mice were genotyped by the Garvan Molecular Genetics (GMG) facility at the Garvan Institute of Medical Research.

#### Mouse strains

*Stat3^T716M^* and *Stat3^K658N^* mice were produced by CRISPR/*Cas9* gene targeting in mouse embryos, following established molecular and animal husbandry techniques (Yang et al., 2014). Target gene-specific single guide RNAs (sgRNA; 15ng/μl) were microinjected into the nucleus and cytoplasm of mouse zygotes, together with polyadenylated S. pyogenes *Cas9* mRNA (30ng/μl) and a gene-specific 150 base single-stranded, deoxy-oligonucleotide homologous recombination substrate (15ng/μl). Founder mice heterozygous for alleles successfully modified by homologous recombination were back-crossed with syngeneic partners and then inter-crossed to establish the *Stat3^T716M^* or *Stat3^K658N^* mouse lines.

For *Stat3^T716M^*, the sgRNA was produced based on a target site in exon 23 (CAGGTCAATGGTATTGCTGC*<u>AGG</u>* = T716M, PAM italicised and underlined) of *Stat3*. The specific sgRNA was microinjected into the nucleus and cytoplasm of C57BL/6J zygotes. The deoxy-oligonucleotide encoded the T716M (A<u>C</u>G>A<u>TG</u>) substitution and a PAM-inactivating silent mutation in the T717 codon (AC<u>C</u>>AC<u>A</u>). For *Stat3^K658N^*, the sgRNA was produced based on a target site in exon 21 (CATGGATGCGACCAACATCC*<u>TGG</u>* = K658N, PAM italicised and underlined) of *Stat3*. The specific sgRNA was microinjected into the nucleus and cytoplasm of C57BL/6J x FVB/N F1 zygotes. The deoxy-oligonucleotide encoded the K658N (AA<u>G</u>>AA<u>C</u>) substitution and a PAM-inactivating silent mutation in the L666 codon (CT<u>G</u>>CT<u>C</u>).

C57BL/6 JAusb (C57BL/6J), C57BL/6 NCrl, B6.SJL-*Ptprc^a^Pepc^b^* (*Ptprc^a/a^*) and B6.129S7-*Rag1^tm1Mom^*/J (*Rag1^KO/KO^*) mice were purchased from ABR.

HyHEL10-transgenic (SWHEL^)^ mice described previously (Phan et al., 2003) carry a single copy VH10 anti-hen egg lysozyme (HEL) heavy chain variable region coding exon targeted to the endogenous *Igh^b^* allele, and multiple copies of the VH10-κ anti-HEL light chain transgene. SWHEL mice on a CD45.1 congenic C57BL/6 background were also *Rag1^KO/KO^*, thus preventing endogenous Ig variable region gene rearrangements (Mombaerts et al., 1992) so that all B-cells expressed the HyHEL10 BCR. SWHEL.*Rag1^KO/KO^* and *Stat3^T716M^* mice were crossed to obtain mice whose B cells all expressed HyHEL10, and were *Stat3^T716M^* wild-type, heterozygous or homozygous. HEL^3X^ is recombinant hen egg lysozyme protein modified by three amino acid substitutions R21Q, R73E, and D101R in the HyHEL10 binding site (Paus et al., 2006), bound by HyHEL10 at relatively low affinity (1.1 x 10^-7^ M) (Chan et al., 2012). Membrane HEL^3X^ transgenic (mHEL^3X^-Tg) mice, on a C57BL6/J background, express the *Hel^3X^* transgene under control of the human ubiquitin C promoter, resulting in ubiquitous expression of membrane-bound HEL^3X^ on the surface of nucleated and anucleate cells (Chan et al., 2012).

#### Chimeras

To generate “100% chimeras”, age- and sex-matched *Rag1^KO/KO^* mice were irradiated with one dose of 425 Rad from an X-ray source (X-RAD 320 Biological Irradiator, PXI). These recipient mice were then injected with donor bone marrow from *Stat3^T716M^* or *Stat3^K658N^* wild-type, heterozygous or homozygous mutant mice.

To generate mixed chimeras, age- and sex-matched *Rag1^KO/KO^* mice were irradiated with 2 doses of 425 Rad 8 hours apart, injected with a 1:1 mixture of *Ptprc^a/a^* bone marrow from congenic *Ptprc^a^* donor mice with *Ptprc^b/b^* cells from *Stat3^T716M^* wild-type or homozygous mutant mice or independently with *Ptprc^a/b^* or *Ptprc^b/b^* cells from *Stat3^K658N^* wild-type or homozygous mutant mice. The donor bone marrow was depleted of lineage-positive cells by MACS using a cocktail of antibodies (to B220, CD3, CD4, CD8, CD11b, CD11c, CD19, LY-6C, LY-6G, NK1.1, TCRβ) prior to injection. Each recipient mouse received 2-6 x 10^6^ lineage-depleted donor bone marrow cells injected intravenously.

Mixed chimeras for the study of central tolerance were generated as previously described (Burnett et al., 2018). Briefly, non-transgenic or mHEL^3X^-Tg *Ptprc^b/b^* C57BL/6 mice were used as recipients, transferred with a bone marrow mixture constituted of 4 parts SWHEL *Rag1^KO/KO^ Ptprc^a/a^* cells and 1 part mHEL^3X^-Tg or non-transgenic *Ptprc^b/b^* C57BL/6 bone marrow, respectively.

Chimeras were analysed 8-14 weeks after reconstitution.

#### Flow cytometry and cell-sorting

Peripheral blood mononuclear cells (PBMCs) were prepared from total blood collected from individuals with STAT3 GOF syndrome. In mice, single-cell suspensions were prepared from spleen, bone marrow, inguinal lymph nodes, peritoneal cavity and blood. 1-4 x 10^6^ cells in PBS 2% FCS were transferred into appropriate wells of a 96-well U bottom plate. To prevent non-specific antibody binding, cells were incubated with Fc blocking antibody for 20 min at 4°C in the dark.

Cells were then incubated with antibodies for 30 min, on ice and in the dark. To fix cells, they were incubated in 10% formalin (Sigma-Aldrich) for 15 min at 4°C, and washed and resuspended in PBS 2% FCS. To stain for intracellular nuclear proteins, cells were fixed and permeabilised using the manufacturer’s instructions and the eBioscience Transcription Factor Staining kit. Stained single-cell suspensions were acquired on the BD LSRFortessa^TM^.

Notably, murine CD19^+^ B cell populations that can express low levels of cell-surface CD21 and CD23 were excluded from the flow analysis of mouse splenic mature CD21^low^ CD23^low^ B cells, including B1a (B220^int^ CD5^high^ CD43^high^ CD23^low^), germinal centre (IgD^-^ CD38^low^ CD95^+^) and immature (CD93^+^) B cells.

Where relevant, following extracellular antibody staining, immune populations were sorted by fluorescence-activated cell sorting (FACS) on a FACS Aria III (BD Biosciences), or by magnetic-activated cell sorting (MACS) on Miltenyi Biotec LS columns, using the procedure indicated by the manufacturer.

#### Anti-mouse antibodies used for flow cytometric analyses

**Anti-mouse antibodies used for flow cytometric analyses.**
| **Cat number** | **Antibody/dye** | **Fluorochrome** | **Company** | **Clone** |
| --- | --- | --- | --- | --- |
| A1310 | 7AAD | N/A | Invitrogen | N/A |
| 17-0051-81 | CD5 | APC | Thermo Fischer | 53-7.3 |
| 557396 | CD11b | FITC | BioLegend | M1/70 |
| 101208 | CD11b | PE | BioLegend | M1/70 |
| 45-0114-82 | CD11c | PerCP/Cy5.5 | eBioscience | N418 |
| 115546 | CD19 | BV510 | BioLegend | 6D5 |
| 115539 | CD19 | BV605 | BioLegend | 6D5 |
| 150405 | CD20 | BV421 | BioLegend | SA275A11 |
| 553818 | CD21/35 | FITC | BD Biosciences | 7G6 |
| 552957 | CD21/35 | PE | BD Biosciences | 7G6 |
| 126111 | CD22 | PE | BioLegend | OX-97 |
| 101614 | CD23 | PE/Cy7 | BioLegend | B3B4 |
| 563987 | CD23 | BV711 | BD Biosciences | B3B4 |
| 101820 | CD24 | Pacific Blue | BD Biosciences | M1/69 |
| 562768 | CD38 | BV421 | BD Biosciences | Ab90 |
| 553270 | CD43 | FITC | BD Biosciences | S7 |
| 564574 | CD45.1 | BUV737 | BD Biosciences | A20 |
| 110716 |  | APC Cy7 | Biolegend | A20 |
| 110728 |  | PerCP Cy5.5 | Biolegend | A20 |
| 560696 | CD45.2 | PE Cy7 | BD Biosciences | 104 |
| 564616 |  | BUV395 | BD Biosciences | 104 |
| 109824 |  | APC Cy7 | BioLegend | 104 |
| 564449 | CD45R/B220 | BUV737 | BD Biosciences | RA3-6B2 |
| 104508 | CD69 | PE | BioLegend | H1.2F3 |
| 740541 | CD72 | BV650 | BD Biosciences | K10.6 |
| 740877 | CD86 | BV786 | BD Biosciences | GL1 |
| 17-5892-83 | CD93 | APC | eBioscience | AA4.1 |
| 557653 | CD95 | PE Cy7 | BD Biosciences | Jo2 |
| 129804 | CD196 (CCR6) | PE | BioLegend | 29-2L17 |
| 129816 |  | PE Cy7 | BioLegend | 29-2L17 |
| 565988 | IgD | BUV395 | BD Biosciences | 11-26c.2a |
| 559750 | Ig, $\kappa$ light chain | Biotin | BD Biosciences | 187.1 |
| 407308 | Ig, $\lambda$ light chain | PE | BioLegend | RML-42 |
| 406515 | IgM | APC/Cy7 | BioLegend | RMM-1 |
| 405229 | Streptavidin | BV605 | BioLegend | N/A |
| 644814 | T-bet | APC | BioLegend | 4B10 |

Normal rat serum was purchased from Sigma and normal mouse serum from Jackson Immunoresearch. HyHEL9 antibody was derived from a hybridoma, purified and conjugated using the AF647 labelling kit (Invitrogen).

#### *In vitro* B cell stimulation

Single cell suspensions of *Stat3* wild-type or mutant mouse spleens were incubated with anti- CD23 biotin followed by MACS positive enrichment for CD23^+^ cells. 2x10^6^ CD23^+^ cells were plated per well in 200 μL complete RPMI (cRPMI), or cRPMI supplemented with 5 μg/mL anti-mouse IgM F(ab’)2 and 5 μg/mL anti-mouse CD40. Cells were cultured for 3 days, washed and stained for flow cytometric analysis.

#### Intracellular calcium measurements

Single cell suspensions of *Stat3* wild-type or mutant splenocytes at a concentration of 2 x 10^6^ cells per mL in cRPMI were pre-loaded with Indo-1-AM (2 μg/mL) for 30 minutes at 37°C. The cells were then acquired on the BD LSR Fortessa^TM^, and at 90 seconds acquisition were exposed to F(ab’)2 goat anti-mouse IgM (60 μg/mL), followed by addition at 300 second of the calcium ionophore Ionomycin, to act as a positive control. Calcium-dependent changes in Indo-1-AM emission at 400 nm were compared between cell populations and between mice using the FlowJo kinetic function.

#### BCR deep sequencing

RNA was extracted from B cell populations bulk-sorted by FACS using the AllPrep DNA/RNA Mini Kit (Qiagen #80204), and reverse transcribed to cDNA using the Smart-Seq2 protocol (Picelli et al., 2014). 10 cycles of PCR amplification were performed and the TSO oligo was modified to incorporate a 10 bp unique molecular identifier (UMI). Following purification of PCR products with magnetic AMPure XP beads (Agencourt), a BCR-specific PCR was performed with a forward primer (ADP_fwd) targeting the 5’ incorporated TSO oligo and a reverse primer targeting the constant region of the mouse IGH chain.

PCR was performed using the KAPA HiFi HotStart Ready Mix (Kappa Biosystems), under the following conditions: 98 °C 45s; 30 cycles: 98 °C 15s, 60 °C 30s, 72 °C 30s; 72 °C 1min. The PCR products were purified using AMPure beads and complete adaptor sequences and sample barcodes were added using the primers from the Illumina Nextera Index Kit (Illumina). 5 cycles of PCR amplification were performed using the Q5 High-Fidelity DNA Polymerase kit (New England BioLabs): 72 3min; 98 °C 30s; 5 cycles: 98 °C 10s, 63 °C 30s, 70 °C 3min. Following a second round of magnetic bead purification, the BCR libraries were quantified using the Qubit 4 fluorometer (Invitrogen) and pooled at equal concentration for sequencing on an Illumina MiSeq 250 bp paired-end run.

Sequencing was performed on the Illumina MiSeq platform using the MiSeq Reagent Kit v3 with a read length mode of 2 x 300bp. Libraries were sequenced to ∼1 million reads per sample.

#### Single-cell RNA sequencing using the 10X platform

To assess the effects of STAT3 GOF on gene expression in B cells, we sorted splenic mature follicular (*n* = 2 donors per genotype) and CD21^low^ CD23^low^ (*n* = 4 donors per genotype) B cells from *Stat3^+/+^* (WT) and *Stat3^K658N/K658N^* (HOM) mice. Mouse B cells were bulk-sorted into Eppendorf tubes containing cold sterile PBS 10% FCS and incubated for 20 min at 4°C with TotalSeq^TM^ DNA-barcoded anti-mouse ‘Hashing’ antibodies (BioLegend) at a 1/100 final dilution. The TotalSeq^TM^ antibodies contain a mixture of two monoclonal antibodies, both conjugated to the same DNA oligonucleotide, that are specific against mouse CD45 and MHC class I haplotypes – and thus stain all leukocytes from C57BL/6 mice.

During the incubation, cells were transferred into a 96-well round bottom plate, on ice.

Following incubation, cells were washed three times in cold PBS 2% FCS and the hashed populations pooled into mixtures for single-cell RNA sequencing using the 10X Genomics platform. The Garvan-Weizmann Centre for Cellular Genomics (GWCCG) performed the 10X capture, and sequencing of resulting cDNA samples, as an in-house commercial service, using the Chromium Single-Cell v2 3’ Kits (10X Genomics). A total of 5,000 to 12,000 cells were captured per reaction.

RNA libraries were sequenced on an Illumina NovaSeq 6000 (NovaSeq Control Software v 1.6.0 / Real Time Analysis v3.4.4) using a NovaSeq S4 230 cycles kit (Illumina, 20447086) as follows: 28bp (Read 1), 91bp (Read 2) and 8bp (Index). HASHing libraries were sequenced on an Illumina NextSeq 500/550 (NextSeq Control Software v 2.2.0.4 / Real Time Analysis 2.4.11) using a NextSeq 60 cycles kit (Illumina, 20456719) as follows: 28bp (Read 1), 24bp (Read 2) and 8bp (Index). Sequencing generated raw data files in binary base call (BCL) format. These files were demultiplexed and converted to FASTQ using Illumina Conversion Software (bcl2fastq v2.19.0.316). Alignment, filtering, barcode counting and UMI counting were performed using the Cell Ranger Single Cell Software v3.1.0 (10X Genomics). Reads were aligned to the mm10-3.0.0 (release 84) mouse reference genomes. Raw count matrices were exported and filtered using the EmptyDrops package in R (Lun et al., 2019).

#### DNA-barcoded anti-mouse Hashing antibodies

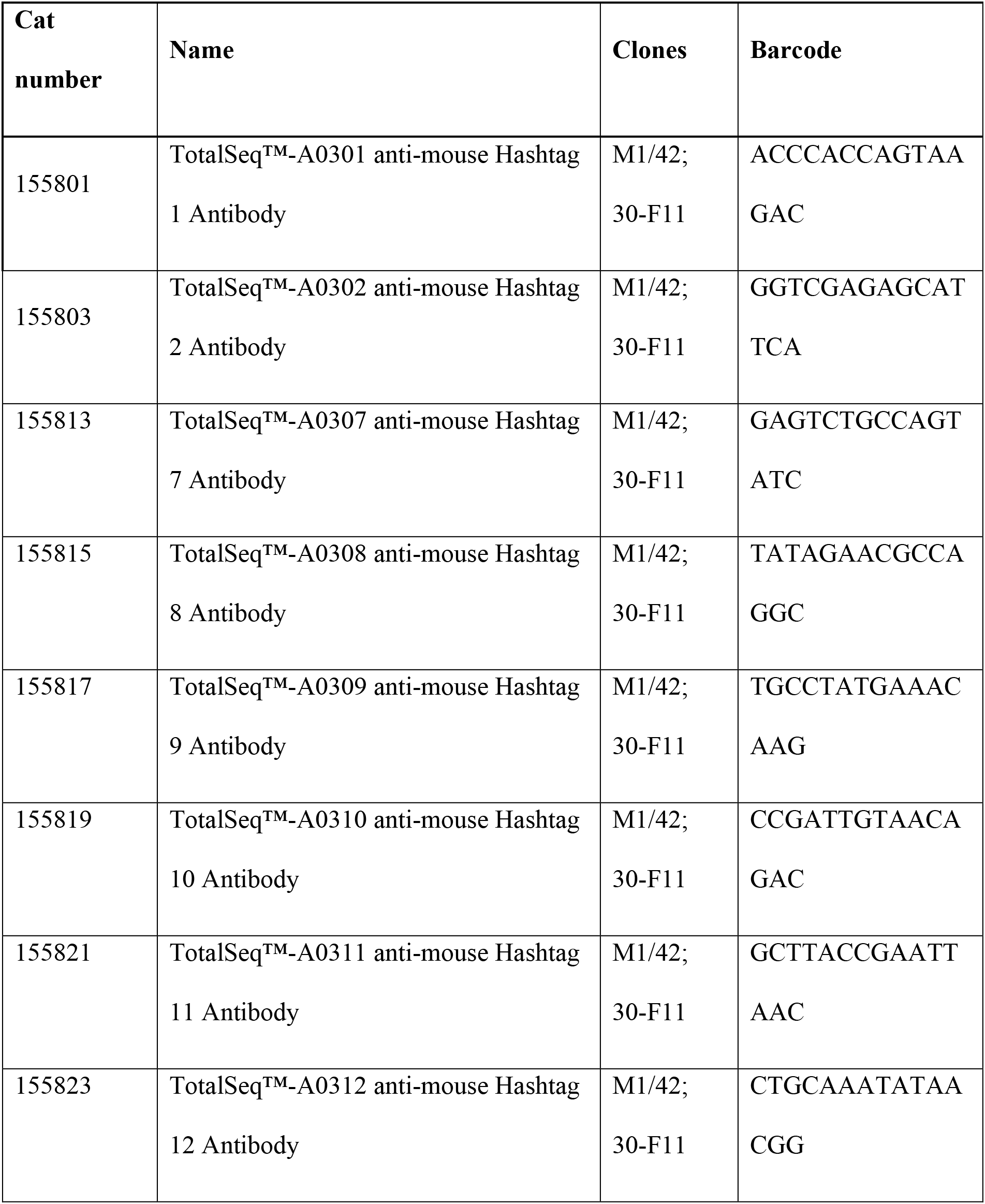

### QUANTIFICATION AND STATISTICAL ANALYSIS

Statistical analyses of flow cytometric experiments were performed using the GraphPad Prism 6 software (GraphPad, San Diego, USA). A one-tailed unpaired Student’s *t*-test with Welch’s correction was used for comparisons between two normally distributed groups. An unpaired student’s *t*-test, corrected for multiple comparisons using the Holm-Sidak method, was used for comparisons of more than two groups. Differences between paired measurements were analysed by paired *t*-test. In all graphs presented, unless otherwise stated, the error bars represent the mean and standard deviation. * p < 0.05, ** p < 0.01, *** p < 0.001.

For the 10X analysis, cells were excluded if the library size or number of expressed genes fell below 2 median absolute deviations, or if mitochondrial reads accounted for more than 20% of total reads. Cell-wise gene expression counts were normalized and recovered using SAVER (Huang et al., 2018) with default values, and differentially expressed genes (DEGs) were identified using limma (Ritchie et al., 2015) on the log-transformed recovered counts. Where appropriate, fold changes and *p*-values were reported after correcting for the *Ptprc* genotype effect during the linear modelling process, through a set of post-hoc contrasts. Bonferroni correction was applied to each set of *p*-values. DEGs were defined as having a family-wise error rate (FWER) < 0.05. UMAPs were generated using Seurat (Satija et al., 2015).

BCR deep sequencing libraries were de-multiplexed by their sample indices using the Illumina FASTQ generation workflow. Paired-end reads were merged using FLASH (Magoc and Salzberg, 2011). Quality filtering, UMI extraction, primer trimming and de-replication were performed using pRESTO (Vander Heiden et al., 2014). The resulting FASTA formatted sequence datasets were aligned against the germline reference directory for the locus and species obtained from IMGT [http://www.imgt.org/] using a local installation of IgBLAST (Ye et al., 2013).

IgBLAST alignment determined the V, D and J gene segments that contributed to each rearrangement, any nucleotide insertions or deletions when the segments were joined, the FR and CDRs, and for the BCR libraries, the nucleotide substitutions from SHM.

## SUPPLEMENTAL INFORMATION TITLES AND LEGENDS

**Figure S1. [Splenic immature B cell populations], related to Figure 1**.

**A,B,** Representative flow cytometric analysis **(A)** and frequencies **(B)** of splenic CD19^+^ B cells, CD93^neg^ mature, CD93^neg^ immature CD23^neg^ T1, CD23^pos^ IgM^high^ T2 and CD23^pos^ IgM^low^ T3 B cells, in *Stat3^+/+^* (grey fill), *Stat3^K658N/+^* (orange fill) or *Stat3^K658N/K658N^* (red fill) mice. **C,D** Representative flow cytometric analysis **(C)** and frequencies **(D)** of splenic B cells, mature, immature T1, T2 and T3 B cells, in *Stat3^+/+^* (grey fill), *Stat3^T716M/+^* (light blue fill) or *Stat3^T716M/T716M^* (dark blue fill) mice. Data representative of *n* > 3 experiments with *n* > 4 mice per group. Comparisons by *t*-test, corrected for multiple comparisons using the Holm-Sidak method. * p < 0.05; ** p < 0.01; *** p < 0.001.

**Figure S2. [*Stat3^K658N/K658N^* causes accumulation of CD21^low^ age-associated-like B cells in circulation and in the lymph nodes], related to Figure 1**.

**A,** Percentage of bone marrow CD23^+^, CD23^-^ CD21^+^ or CD23^-^ CD21^-^ mature recirculating B cells. Symbols denote values from individual *Stat3^+/+^* (grey fill), *Stat3^T716M/+^* (light blue fill) or *Stat3^T716M/T716M^* (dark blue fill) mice, or from *Stat3^+/+^* (grey fill), *Stat3^K658N/+^* (orange fill) or *Stat3^K658N/K658N^* (red fill) mice. **B, C,** Representative flow cytometric analysis of CD21 and CD23 expression by mature CD19^+^ B220^+^ CD93^-^ CD95^-^ B cells, and percentage or total number in the blood (**B**) and inguinal lymph nodes (**C**) of mature CD21^low^ CD23^low^ B cells, in mice of the indicated genotypes. **D,** *Rag1^KO/KO^* mice were transplanted with bone marrow from *Stat3^+/+^ Ptprc^a/a^* (black fill) donor mice in a 1:1 mixture with bone marrow from a *Stat3^+/+^* (grey fill) or *Stat3^K658N/K658N^* (red fill) *Ptprc^a/b^* donor mouse. Solid lines link cells within one chimeric mouse. Plots show mean fluorescence intensity (MFI) following flow cytometric analysis of cell-surface CD21 (left) or CD23 (right) expression by splenic CD93^+^ CD23^-^ (T1), CD93^+^ CD23^+^ IgM^high^ T2, CD93^+^ CD23^+^ IgM^low^ T3, CD93^-^ CD23^+^ follicular (FO), CD93^-^ CD23^low^ CD21^+^ marginal zone (MZ) and CD93^-^ CD23^low^ CD21^low^ B cells of the indicated genotypes. **(A-C)** Data representative of *n* > 2 experiments with *n* > 4 mice per group. Comparisons made by *t*-test, corrected for multiple comparisons using the Holm-Sidak method. * p < 0.05; ** p < 0.01; *** p < 0.001. **(D)** Data representative of *n* = 2 experiments with *n* > 4 mice per group. Comparisons made by paired *t*-test.

**Figure S3. [CD21^low^ CD23^low^ B cells have a gene expression profile similar to previously published CD21^low^ CD23^low^ age-associated B cell populations], related to Figure 4**.

Comparison of sorted CD21^low^ CD23^low^ and follicular B cells (*Stat3^K658N^* HOM and WT) by single-cell RNA sequencing. **A,** Comparison by gene set enrichment analysis, analysing GSE28887 comprising a previously published set (Rubtsov et al., 2011) of mRNAs decreased (upper panel) or increased (lower panel) in age-associated B cells (ABC; sorted as B220^+^ CD19^+^ CD11b^+^ CD11c^+^) compared to follicular B cells (B220^+^ CD19^+^CD11b^-^ CD1d^jnt^ CD21^int^) sorted from elderly wild-type C57BL/6 mice. **B,** Comparison as volcano plot of log2 expression fold-change (log2FC) versus moderated *t*-statistic for differentially expressed genes between CD21^low^ CD23^low^ and follicular B cells. Circles denote all genes with log2FC≠0 and FWER<0.05. Purple or green text denote increased or decreased genes, respectively, previously reported as differentially expressed by “age-associated”, “CD21^low^”, “double-negative”, “CD11c^+^” or “atypical memory” B cells relative to mature naïve follicular or memory B cells (Charles et al., 2011, Jenks et al., 2018, Lau et al., 2017, Rakhmanov et al., 2009, Rubtsov et al., 2011, Russell Knode et al., 2017, Scharer et al., 2019). Blue denotes the two genes (*Fcer2a* and *Cr2*) whose protein products (CD23 and CD21) were used to sort CD21^low^ CD23^low^ mature B cells from CD21^med^ CD23^+^ follicular B cells. **C, D,** Venn diagrams of intersection between genes binding phosphorylated STAT3 (pSTAT3) by chromatin immunoprecipitation sequencing (ChIP-seq) of BCR-stimulated primary human peripheral naïve B cells and of TMD8 cells (Lu et al., 2019) (black circles) and genes with significantly (FWER<0.05) increased (red circle) or decreased (green circle) expression in HOM relative to WT CD21^low^ CD23^low^ B cells. FWER of intersection by Fisher exact test: increased genes, p = 3.92 x 10^-35^ (odds ratio 2.06); decreased genes, p = 1.15 x 10^-14^ (odds ratio 1.39).

**Figure S4. [STAT3 gain-of-function causes a cell-intrinsic accumulation of CCR6^high^ and CD11b^+^ CD21^low^ CD23^low^ B cells], related to Figure 4**.

(**A-D**) Mixed chimeras were generated by transplanting *Rag1^KO/KO^* mice with bone marrow from a *Ptprc^a/a^ Stat3^+/+^* (CD45.1^+^; black fill) donor, in a 1:1 mixture with bone marrow from a *Ptprc^a/b^* (CD45.2^+^) donor that was *Stat3^+/+^* (grey fill), *Stat3^K658N/K658N^* (red fill). The same workflow was performed independently using *Ptprc^b/b^ Stat3^+/+^* and *Stat3^T716M/T716M^* donors. Black lines connect cells from the same chimeric mouse. **A**, Representative flow cytometric analysis of CCR6 versus CD23 cell-surface expression, by B220^+^ CD93^-^ mature B cells of the indicated genotypes in mixed chimeras that received *Ptprc^a/b^ Stat3^K658N^* (top) or *Ptprc^b/b^ Stat3^T716M^* (bottom) wild-type or mutant bone marrow. **B**, Mean fluorescence intensity (MFI) of CCR6 cell-surface expression following flow cytometric analysis of splenic B cells of the indicated genotypes. **C**, Representative flow cytometric analysis of CD11b (top row) or CD11c (bottom row) versus CD23 cell-surface expression, by B220^+^ CD93^-^ mature B cells of the indicated genotypes in mixed chimeras that received *Stat3^K658N^* wild-type or homozygous mutant *Ptprc^a/b^* bone marrow. **D**, Mean fluorescence intensity (MFI) of CD11b cell-surface expression following flow cytometric analysis of splenic B cells of the indicated genotypes. **E**, Representative plots of CD21^low^ CD23^low^ B cells showing subsets with a CD11b^+^ CD11c^-^, CD11b^+^ CD11c^+^ or CD11b^-^ CD11c^+^ phenotype, and the number of each subset per spleen or per μL of blood in individual *Stat3^+/+^* (grey fill), *Stat3^K658N/+^* (orange fill) or *Stat3^K658N/K658N^* (red fill) mice. **F**, Percentage of splenic B cells that were mature CD23^-^ CD11b^+^ cells in *Stat3^+/+^* (grey fill), *Stat3^T716M/+^* (light blue fill) or *Stat3^T716M/T716M^* (dark blue fill) mice. (**A-D**) Data are representative of *n* = 2 independent experiments with *n* > 4 recipients per group.

Comparisons within each recipient mouse made by paired t-test. Comparisons between mice made by one-way ANOVA followed by multiple comparison post-tests. (**E,F**) Data are representative of *n* = 3 experiments with *n* > 3 mice per group. Comparisons made by t-test, corrected for multiple comparisons using the Holm-Sidak method. * p < 0.05; ** p < 0.01; *** p < 0.001.

**Figure S5. [Altered cell-surface IgM and IgD expression on B cell populations from *Stat3^K658N/K658N^* mice], related to Figure 5**.

**A**, Representative histogram overlays of cell-surface CD22 expression by B220^+^ CD19^+^ CD95^-^ CD93^-^ CD21^low^ CD23^low^ B cells of the indicated genotypes, in mixed chimeras that received: *Ptprc^b/b^ Stat3^+/+^* bone marrow in a 1:1 mixture with *Ptprc^a/b^ Stat3^+/+^* or *Stat3^K658N/K658N^* bone marrow (top) or *Ptprc^a/a^ Stat3^+/+^* bone marrow in a 1:1 mixture with *Ptprc^b/b^ Stat3^+/+^* or *Stat3^T716M/T716M^* bone marrow (bottom). Data are representative of *n* > 2 experiments for both strains, with *n* > 4 recipients per donor genotype. **B**, Representative histogram overlays of cell-surface IgD (left) and IgM (right) expression on *Stat3^K658N/K658N^* CD21^low^ CD23^low^ B cells (red line and fill) compared to *Stat3^+/+^* CD21^low^ CD23^low^ B cells (solid black line) and *Stat3^+/+^* follicular B cells (dotted black line). Data are representative of *n* > 3 experiments with *n* > 4 mice per group. **C**, Mean fluorescence intensity (MFI) following fluorescent antibody staining for cell-surface IgM (top) or IgD (bottom) expression by splenic B cell populations in *Stat3^+/+^* (grey fill), *Stat3^K658N/+^* (orange fill) or *Stat3^K658N/K658N^* (red fill) mice 7-12 weeks old. Data are representative of *n* > 3 experiments with *n* > 4 mice per group. Comparisons made by t-test, corrected for multiple comparisons using the Holm-Sidak method. * p < 0.05; ** p < 0.01; *** p < 0.001.

**Table S1. [**Genes differentially expressed in CD21^low^ CD23^low^ relative to naïve follicular mature B cells with wild-type (WT) or K658N homozygous (HOM) *Stat3*], related to Figure 5.

**Table S2. [**Results of gene set enrichment analysis comparing CD21^low^ CD23^low^ relative to follicular B cells for sets of genes up-regulated in B220^+^ CD93^-^ CD43^-^ CD21^-^ CD23^-^ (GSE81650) or B220^+^ CD19^+^ CD11b^+^ (GSE28887) age-associated B cells relative to follicular B cells], related to Figure 5.

**Table S3. [**Genes differentially expressed in *Stat3^K658N/K658N^* relative to *Stat3^+/+^* CD21^low^ CD23^low^ B cells], related to Figures 5 & 6.

**Table S4. [**Intersection of genes differentially expressed in *Stat3^K658N/K658N^* relative to *Stat3^+/+^* CD21^low^ CD23^low^ B cells, and also published by Lu *et al*. as STAT3- or pSTAT3-bound following ChIP-seq of human peripheral B cells and B lymphoma cell lines], related to Figures 5 & 6.

**Table S5. [**Intersection of genes differentially expressed in *Stat3^K658N/K658N^* relative to *Stat3^+/+^* CD21^low^ CD23^low^ B cells, at the leading edge of the KEGG or GO BCR signalling gene sets and pSTAT3-bound in human peripheral B cells and B lymphoma cell lines], related to Figure 6.

