## Supplemental Figures and Legends for "Overactive STAT3 drives accumulation of disease-associated CD21^low^ B cells"

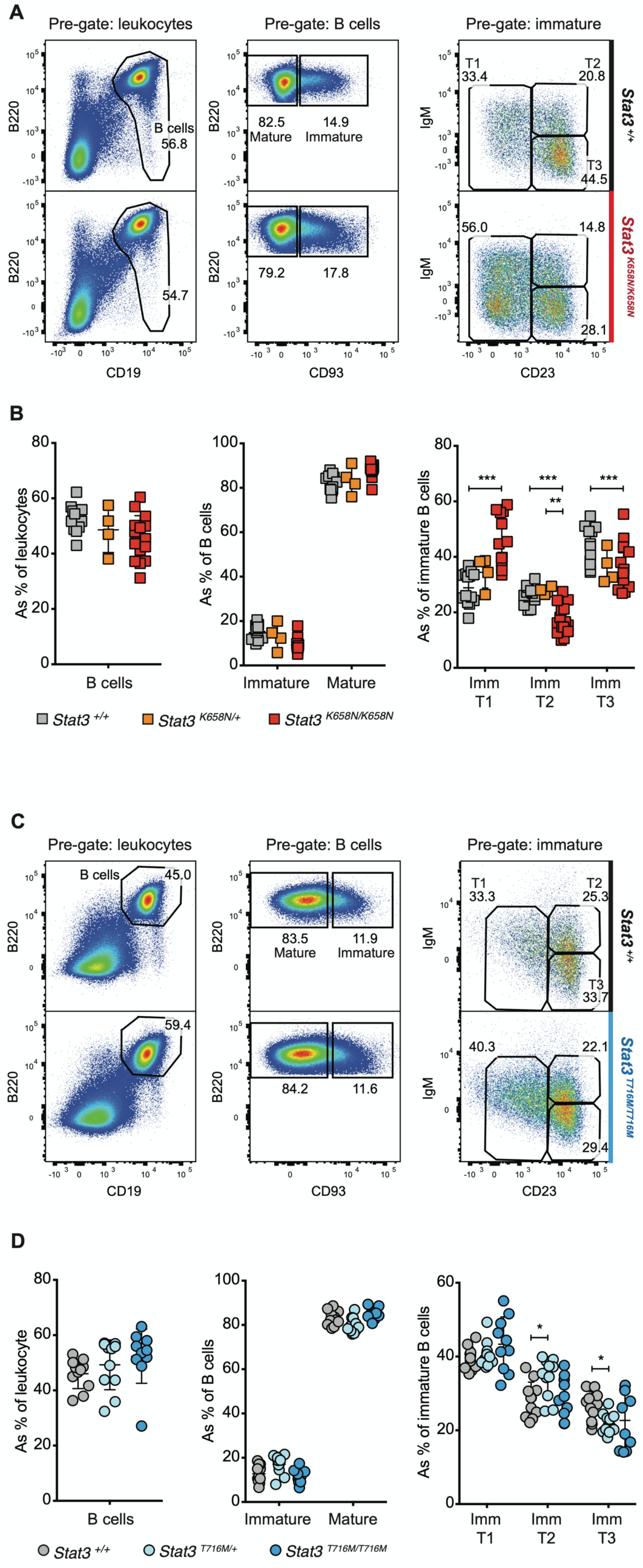


**Figure S1. [Splenic immature B cell populations], related to Figure 1.**

**A,B**, Representative flow cytometric analysis (**A**) and frequencies (**B**) of splenic CD19⁺ B cells, CD93$^{neg}$ mature, CD93$^{neg}$ immature CD23$^{neg}$ T1, CD23$^{pos}$ IgM$^{high}$ T2 and CD23$^{pos}$ IgM$^{low}$ T3 B cells, in *Stat3*$^{+/+}$ (grey fill), *Stat3*$^{K658N/+}$ (orange fill) or *Stat3*$^{K658N/K658N}$ (red fill) mice. **C,D** Representative flow cytometric analysis (**C**) and frequencies (**D**) of splenic B cells, mature, immature T1, T2 and T3 B cells, in *Stat3*$^{+/+}$ (grey fill), *Stat3*$^{T716M/+}$ (light blue fill) or *Stat3*$^{T716M/T716M}$ (dark blue fill) mice. Data representative of $n > 3$ experiments with $n > 4$ mice per group. Comparisons by *t*-test, corrected for multiple comparisons using the Holm-Sidak method. * $p < 0.05$; ** $p < 0.01$; *** $p < 0.001$.

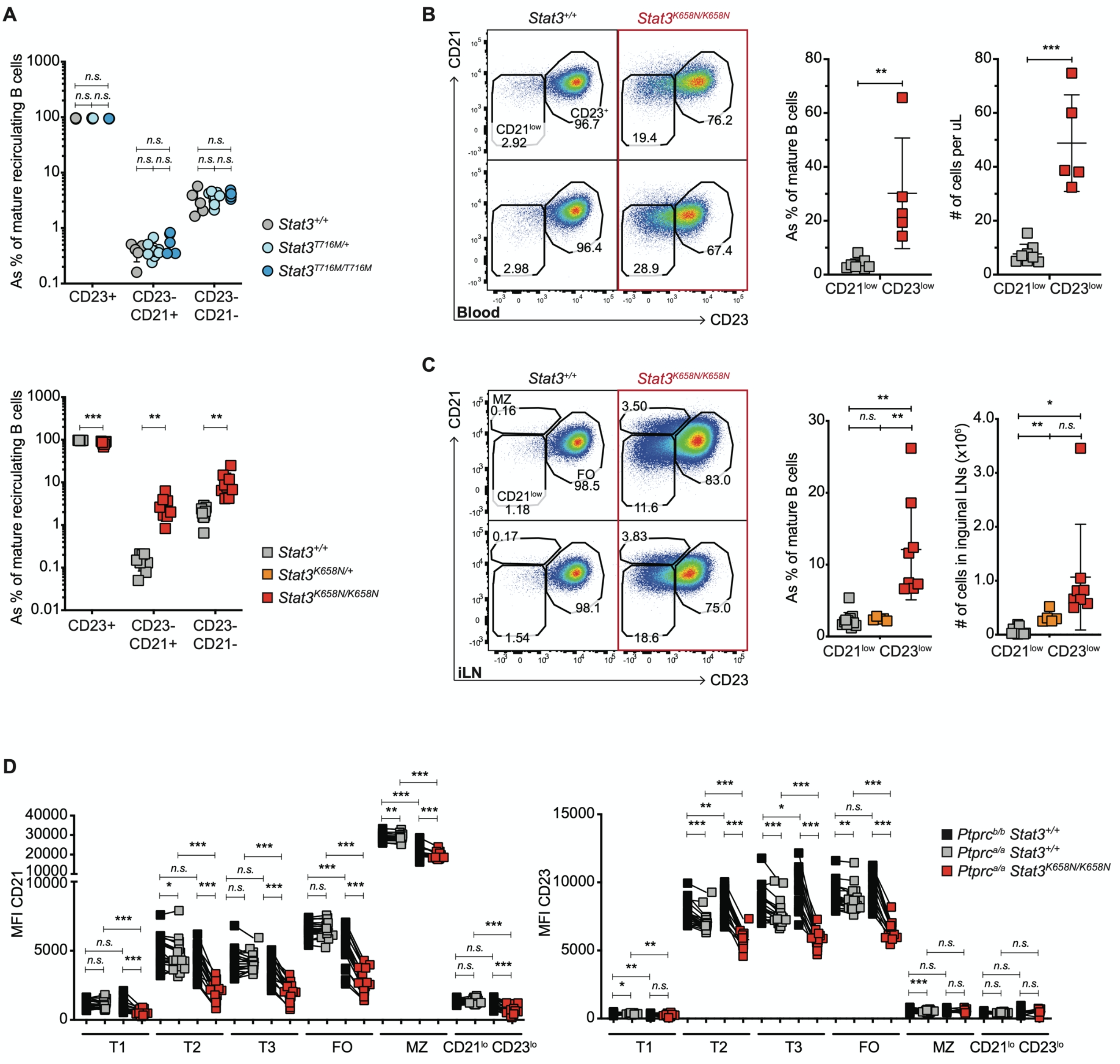


**Figure S2. [*Stat3*$^{K658N/K658N}$ causes accumulation of CD21$^{low}$ age-associated-like B cells in circulation and in the lymph nodes], related to Figure 1.**

**A**, Percentage of bone marrow CD23$^{+}$, CD23$^{-}$ CD21$^{+}$ or CD23$^{-}$ CD21$^{-}$ mature recirculating B cells. Symbols denote values from individual *Stat3*$^{+/+}$ (grey fill), *Stat3*$^{T716M/+}$ (light blue fill) or

*Stat3*$^{T716M/T716M}$ (dark blue fill) mice, or from *Stat3*$^{+/+}$ (grey fill), *Stat3*$^{K658N/+}$ (orange fill) or *Stat3*$^{K658N/K658N}$ (red fill) mice. **B, C,** Representative flow cytometric analysis of CD21 and CD23 expression by mature CD19$^{+}$ B220$^{+}$ CD93$^{-}$ CD95$^{-}$ B cells, and percentage or total number in the blood (**B**) and inguinal lymph nodes (**C**) of mature CD21$^{low}$ CD23$^{low}$ B cells, in mice of the indicated genotypes. **D,** *Rag1*$^{KO/KO}$ mice were transplanted with bone marrow from *Stat3*$^{+/+}$ *Ptprc*$^{a/a}$ (black fill) donor mice in a 1:1 mixture with bone marrow from a *Stat3*$^{+/+}$ (grey fill) or *Stat3*$^{K658N/K658N}$ (red fill) *Ptprc*$^{a/b}$ donor mouse. Solid lines link cells within one chimeric mouse. Plots show mean fluorescence intensity (MFI) following flow cytometric analysis of cell-surface CD21 (left) or CD23 (right) expression by splenic CD93$^{+}$ CD23$^{-}$ (T1), CD93$^{+}$ CD23$^{+}$ IgM$^{high}$ T2, CD93$^{+}$ CD23$^{+}$ IgM$^{low}$ T3, CD93$^{-}$ CD23$^{+}$ follicular (FO), CD93$^{-}$ CD23$^{low}$ CD21$^{+}$ marginal zone (MZ) and CD93$^{-}$ CD23$^{low}$ CD21$^{low}$ B cells of the indicated genotypes. **(A-C)** Data representative of $n > 2$ experiments with $n > 4$ mice per group. Comparisons made by *t*-test, corrected for multiple comparisons using the Holm-Sidak method. * $p < 0.05$; ** $p < 0.01$; *** $p < 0.001$. **(D)** Data representative of $n = 2$ experiments with $n > 4$ mice per group. Comparisons made by paired *t*-test.

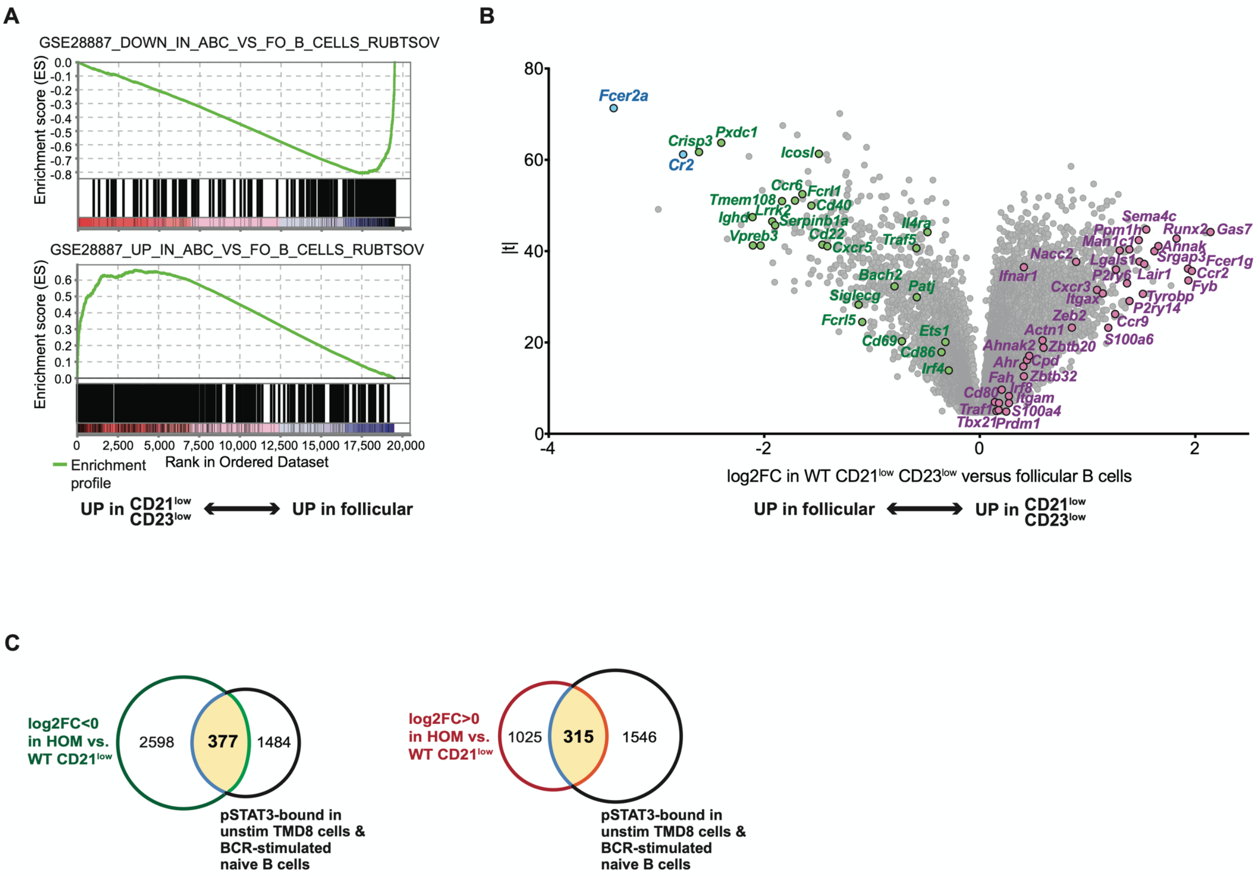

**Figure S3. [CD21$^{low}$ CD23$^{low}$ B cells have a gene expression profile similar to previously published CD21$^{low}$ CD23$^{low}$ age-associated B cell populations], related to Figure 4.**

Comparison of sorted CD21$^{low}$ CD23$^{low}$ and follicular B cells (*Stat3$^{K658N}$* HOM and WT) by single-cell RNA sequencing. **A,** Comparison by gene set enrichment analysis, analysing GSE28887 comprising a previously published set (Rubtsov et al., 2011) of mRNAs decreased (upper panel) or increased (lower panel) in age-associated B cells (ABC; sorted as B220$^{+}$ CD19$^{+}$ CD11b$^{+}$ CD11c$^{+}$) compared to follicular B cells (B220$^{+}$ CD19$^{+}$CD11b$^{-}$ CD1d$^{int}$ CD21$^{int}$) sorted from elderly wild-type C57BL/6 mice. **B,** Comparison as volcano plot of log2 expression fold-change (log2FC) versus moderated *t*-statistic for differentially expressed genes between CD21$^{low}$ CD23$^{low}$ and follicular B cells. Circles denote all genes with log2FC≠0 and FWER<0.05. Purple or green text denote increased or decreased genes, respectively, previously reported as differentially expressed by "age-associated", "CD21$^{low}$", "double-negative", "CD11c$^{+}$" or "atypical memory" B cells relative to mature naïve follicular or memory B cells (Charles et al., 2011, Jenks et al., 2018, Lau et al., 2017, Rakhmanov et al., 2009, Rubtsov et al., 2011, Russell Knode et al., 2017, Scharer et al., 2019). Blue denotes the two genes (*Fcer2a* and *Cr2*) whose protein products (CD23 and CD21) were used to sort CD21$^{low}$ CD23$^{low}$ mature B cells from CD21$^{med}$ CD23$^{+}$ follicular B cells. **C, D,** Venn diagrams of intersection between genes binding phosphorylated STAT3 (pSTAT3) by chromatin immunoprecipitation sequencing (ChIP-seq) of BCR-stimulated primary human peripheral naïve B cells and of TMD8 cells (Lu et al., 2019) (black circles) and genes with significantly (FWER<0.05) increased (red circle) or decreased (green circle) expression in HOM relative to WT CD21$^{low}$ CD23$^{low}$ B cells. FWER of intersection by Fisher exact test: increased genes, p = $3.92 \times 10^{-35}$ (odds ratio 2.06); decreased genes, p = $1.15 \times 10^{-14}$ (odds ratio 1.39).

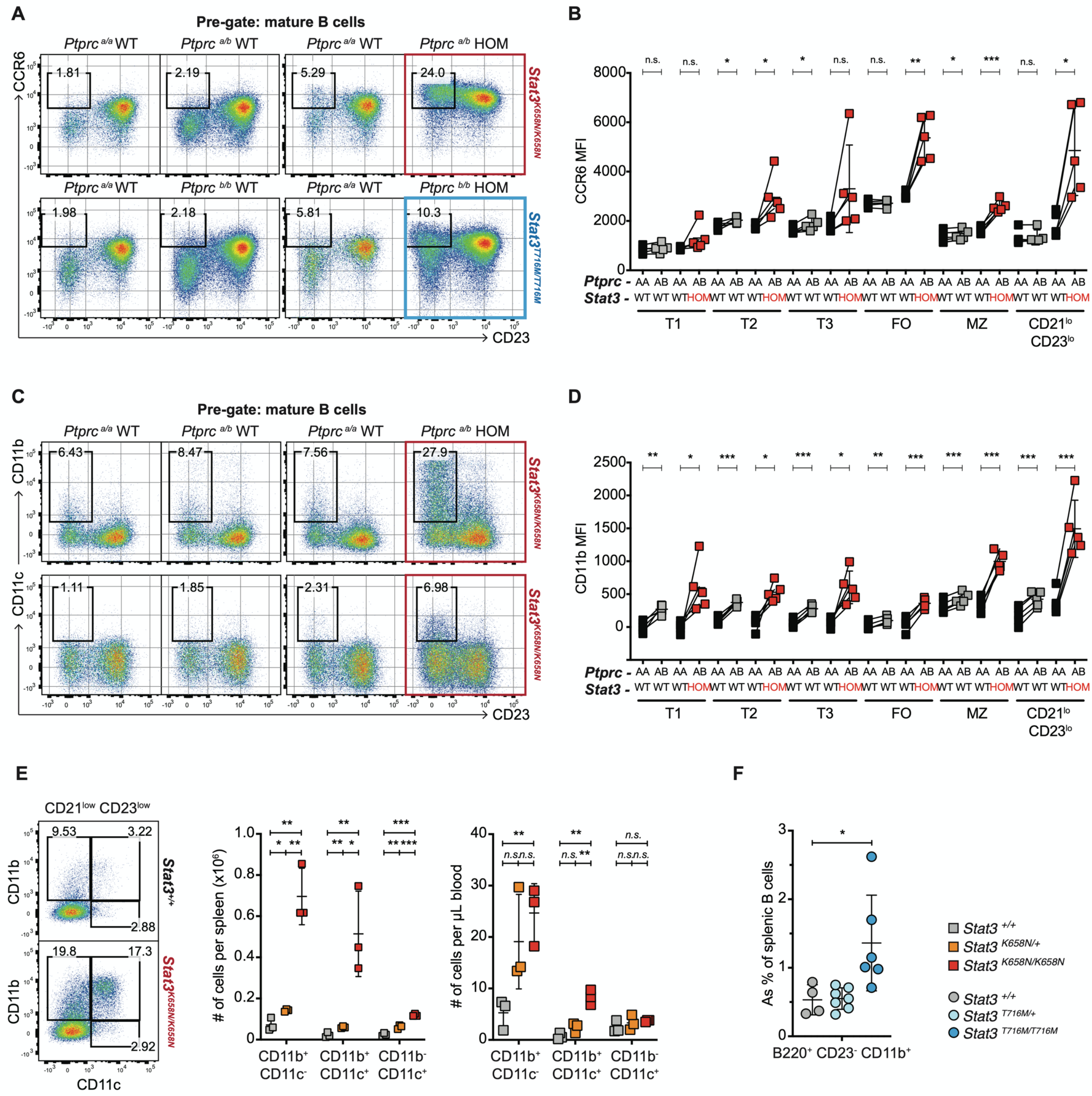


**Figure S4. [STAT3 gain-of-function causes a cell-intrinsic accumulation of CCR6$^{high}$ and CD11b$^{+}$ CD21$^{low}$ CD23$^{low}$ B cells], related to Figure 4.**

(**A-D**) Mixed chimeras were generated by transplanting *Rag1$^{KO/KO}$* mice with bone marrow from a *Ptprc$^{a/a}$ Stat3$^{+/+}$* (CD45.1$^{+}$; black fill) donor, in a 1:1 mixture with bone marrow from a *Ptprc$^{a/b}$* (CD45.2$^{+}$) donor that was *Stat3$^{+/+}$* (grey fill), *Stat3$^{K658N/K658N}$* (red fill). The same workflow was performed independently using *Ptprc$^{b/b}$ Stat3$^{+/+}$* and *Stat3$^{T716M/T716M}$* donors. Black lines connect cells from the same chimeric mouse. **A**, Representative flow cytometric analysis of CCR6 versus CD23 cell-surface expression, by B220$^{+}$ CD93$^{-}$ mature B cells of the indicated genotypes in mixed chimeras that received *Ptprc$^{a/b}$ Stat3$^{K658N}$* (top) or *Ptprc$^{b/b}$ Stat3$^{T716M}$* (bottom) wild-type or mutant bone marrow. **B**, Mean fluorescence intensity (MFI) of CCR6 cell-surface expression following flow cytometric analysis of splenic B cells of the indicated genotypes. **C**, Representative flow cytometric analysis of CD11b (top row) or CD11c (bottom row) versus CD23 cell-surface expression, by B220$^{+}$ CD93$^{-}$ mature B cells of the indicated genotypes in mixed chimeras that received *Stat3*$^{K658N}$ wild-type or homozygous mutant *Ptprc*$^{a/b}$ bone marrow. **D**, Mean fluorescence intensity (MFI) of CD11b cell-surface expression following flow cytometric analysis of splenic B cells of the indicated genotypes. **E**, Representative plots of CD21$^{low}$ CD23$^{low}$ B cells showing subsets with a CD11b$^{+}$ CD11c$^{-}$, CD11b$^{+}$ CD11c$^{+}$ or CD11b$^{-}$ CD11c$^{+}$ phenotype, and the number of each subset per spleen or per µL of blood in individual *Stat3*$^{+/+}$ (grey fill), *Stat3*$^{K658N/+}$ (orange fill) or *Stat3*$^{K658N/K658N}$ (red fill) mice. **F**, Percentage of splenic B cells that were mature CD23$^{-}$ CD11b$^{+}$ cells in *Stat3*$^{+/+}$ (grey fill), *Stat3*$^{T716M/+}$ (light blue fill) or *Stat3*$^{T716M/T716M}$ (dark blue fill) mice. (**A-D**) Data are representative of $n$ = 2 independent experiments with $n$ > 4 recipients per group. Comparisons within each recipient mouse made by paired t-test. Comparisons between mice made by one-way ANOVA followed by multiple comparison post-tests. (**E,F**) Data are representative of $n$ = 3 experiments with $n$ > 3 mice per group. Comparisons made by t-test, corrected for multiple comparisons using the Holm-Sidak method. * $p < 0.05$; ** $p < 0.01$; *** $p < 0.001$.

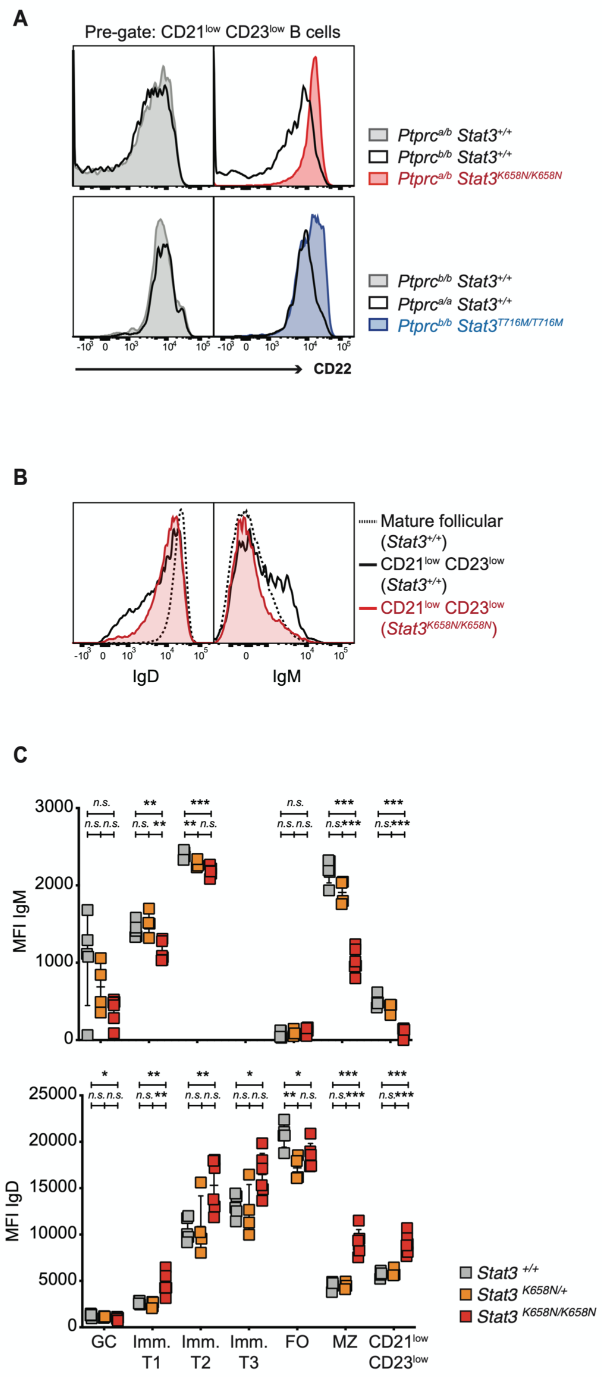


**Figure S5. [Altered cell-surface IgM and IgD expression on B cell populations from *Stat3*$^{K658N/K658N}$ mice], related to Figure 5.**

**A**, Representative histogram overlays of cell-surface CD22 expression by B220$^{+}$ CD19$^{+}$ CD95$^{-}$ CD93$^{-}$ CD21$^{low}$ CD23$^{low}$ B cells of the indicated genotypes, in mixed chimeras that received: *Ptprc*$^{b/b}$ *Stat3*$^{+/+}$ bone marrow in a 1:1 mixture with *Ptprc*$^{a/b}$ *Stat3*$^{+/+}$ or *Stat3*$^{K658N/K658N}$ bone marrow (top) or *Ptprc*$^{a/a}$ *Stat3*$^{+/+}$ bone marrow in a 1:1 mixture with *Ptprc*$^{b/b}$ *Stat3*$^{+/+}$ or *Stat3*$^{T716M/T716M}$ bone marrow (bottom). Data are representative of $n > 2$ experiments for both strains, with $n > 4$ recipients per donor genotype. **B**, Representative histogram overlays of cell-surface IgD (left) and IgM (right) expression on *Stat3*$^{K658N/K658N}$ CD21$^{low}$ CD23$^{low}$ B cells (red line and fill) compared to *Stat3*$^{+/+}$ CD21$^{low}$ CD23$^{low}$ B cells (solid black line) and *Stat3*$^{+/+}$ follicular B cells (dotted black line). Data are representative of $n > 3$ experiments with $n > 4$ mice per group. **C**, Mean fluorescence intensity (MFI) following fluorescent antibody staining for cell-surface IgM (top) or IgD (bottom) expression by splenic B cell populations in *Stat3*$^{+/+}$ (grey fill), *Stat3*$^{K658N/+}$ (orange fill) or *Stat3*$^{K658N/K658N}$ (red fill) mice 7-12 weeks old. Data are representative of $n > 3$ experiments with $n > 4$ mice per group. Comparisons made by t-test, corrected for multiple comparisons using the Holm-Sidak method. * $p < 0.05$; ** $p < 0.01$; *** $p < 0.001$.
