## Supplemental Table 6 for "Overactive STAT3 drives accumulation of disease-associated CD21^low^ B cells"

**KEY RESOURCES TABLE**

| **REAGENT or RESOURCE** | **SOURCE** | **IDENTIFIER** |
| --- | --- | --- |
| **Antibodies** | | |
| 7AAD | Invitrogen | Cat# A1310; RRID: |
| Rat Anti-mouse CD5 APC; Clone 53-7.3 | Thermo Fisher | Cat# 17-0051-81; RRID: AB_469331 |
| Rat Anti-mouse CD11b FITC; Clone M1/70 | BD Biosciences | Cat# 557396; RRID: AB_396679 |
| Rat Anti-mouse CD11b PE; Clone M1/70 | BioLegend | Cat# 101208; RRID: AB_312791 |
| Armenian Hamster Anti-mouse CD11c PerCP-Cy5.5; Clone N418 | Thermo Fisher | Cat# 45-0114-82; RRID: AB_925727 |
| Rat Anti-mouse CD19 BV510; Clone 6D5 | BioLegend | Cat# 115546; RRID: AB_2562137 |
| Rat Anti-mouse CD19 BV605; Clone 6D5 | BioLegend | Cat# 115539; RRID: AB_11203538 |
| Rat Anti-mouse CD21/35 FITC; Clone 7G6 | BD Biosciences | Cat# 553818; RRID: AB_395070 |
| Rat Anti-mouse CD22 PE; Clone OX-97 | BioLegend | Cat# 126111; RRID: AB_ 2561631 |
| Rat Anti-mouse CD23 PE-Cy7; Clone B3B4 | BioLegend | Cat# 101614; RRID: AB_2103036 |
| Rat Anti-mouse CD23 Biotin; Clone B3B4 | BioLegend | Cat# 101604; RRID: AB_312829 |
| Rat Anti-mouse CD24 Pacific Blue; Clone M1/69 | BioLegend | Cat# 101820; RRID: AB_572011 |
| Rat Anti-mouse CD38 BV421; Clone Ab90 | BD Biosciences | Cat# 562768; RRID: AB_2737781 |
| Purified Hamster Anti-Mouse CD40; Clone HM40-3 | BD Biosciences | Cat# 553722; RRID: AB_395007 |
| Rat Anti-mouse CD43 FITC; Clone S7 | BD Biosciences | Cat# 553270; RRID: AB_394747 |
| Mouse Anti-mouse CD45.1 BUV737; Clone A20 | BD Biosciences | Cat# 564574; RRID: AB_2738850 |
| Mouse Anti-mouse CD45.1 APC-Cy7; Clone A20 | Biolegend | Cat# 110716; RRID: AB_313505 |
| Mouse Anti-mouse CD45.1 PerCP-Cy5.5; Clone A20 | Biolegend | Cat# 110728; RRID: AB_893346 |
| Mouse Anti-mouse CD45.2 PE-Cy7; Clone 104 | BD Biosciences | Cat# 560696; RRID: AB_1727494 |
| Mouse Anti-mouse CD45.2 BUV395; Clone 104 | BD Biosciences | Cat# 564616; RRID: AB_2738867 |
| Mouse Anti-mouse CD45.2 APC-Cy7; Clone 104 | BioLegend | Cat# 109824; RRID: AB_830789 |
| Rat Anti-mouse CD45R/B220 BUV737; Clone RA3-6B2 | BD Biosciences | Cat# 612838; Clone: RA3-6B2 |
| Armenian Hamster Anti-mouse CD69 PE; Clone H1.2F3 | BioLegend | Cat# 104508; RRID: AB_313111 |
| Mouse Anti-Mouse CD72 (a, b and d alloantigens); Clone K10.6 | BD Biosciences | Cat# 740541; RRID: AB_2740247 |
| Rat Anti-mouse CD86 BV786; Clone GL1 | BD Biosciences | Cat# 740877; RRID: AB_2740528 |
| Rat Anti-mouse CD93 APC; Clone AA4.1 | Thermo Fisher | Cat# 17-5892-83; RRID: AB_469466 |
| Armenian Hamster Anti-mouse CD95 PE-Cy7; Clone Jo2 | BD Biosciences | Cat# 557653; RRID: AB_396768 |
| Armenian Hamster Anti-mouse CD196 (CCR6) PE-Cy7; Clone 29-2L17 | BioLegend | Cat# 129816; RRID: AB_ 2072798 |
| Rat Anti-mouse IgD BUV395; Clone 11-26c.2a | BD Biosciences | Cat# 565988; RRID: AB_2738723 |
| Rat Anti-mouse Ig, κ light chain Biotin; Clone 187.1 | BD Biosciences | Cat# 559750; RRID: AB_397314 |
| Rat Anti-mouse Ig, λ light chain PE; Clone RML-42 | BioLegend | Cat# 407308; RRID: AB_1027659 |
| Rat Anti-mouse IgM APC-Cy7; Clone RMM-1 | BioLegend | Cat# 406515; RRID: AB_10690815 |
| Affinipure F(ab’)_2_ Fragment Goat anti-Mouse IgM, μ chain specific | Jackson Immunoresearch | Cat# 406515; RRID: AB_ 2338469 |
| Anti-mouse Streptavidin BV605 | BioLegend | Cat# 405229 |
| Mouse Anti-mouse T-bet APC; Clone 4B10 | BioLegend | Cat# 644814; RRID: AB_10901173 |
| Mouse Anti-Human CD19 FITC; Clone 4G7 | BD Biosciences | Cat# 347543 |
| Mouse Anti-Human CD21 BUV563; Clone B-ly4 | BD Biosciences | Cat# 741362; RRID: AB_2870862 |
| Mouse Anti-Human CD20 BUV805; Clone 2H7 | BD Biosciences | Cat# 612905; RRID: AB_2870192 |
| Mouse Anti-Human CD27 BB515; Clone M-T271 | BD Biosciences | Cat# 564642; RRID: AB_2744354 |
| Mouse Anti-Human IgD BV480; Clone IA6-2 | BD Biosciences | Cat# 566138; RRID: AB_2739536 |
| Mouse Anti-Human CD197 (CCR7); Clone 150503 | BD Biosciences | Cat# 561143; RRID: AB_10562031 |
| Mouse Anti-Human CD95 PE; Clone DX2 | BD Biosciences | Cat# 555674; RRID: AB_396027 |
| Mouse Anti-Human CD86 BUV737; Clone 2331 | BD Biosciences | Cat# 612784; RRID: AB_2738804 |
| PerCP/Cy5.5 anti-T-bet antibody; Clone 4B10 | BioLegend | Cat# 644806; RRID: AB_1595488 |
| Indo-1, AM, cell permeant | Thermo Fisher | Cat# I1226 |
| TotalSeq™-A0301 anti-mouse Hashtag 1 Antibody; Clones M1/42, 30-F11 | BioLegend | Cat# 155801; RRID: AB_2750032 |
| TotalSeq™-A0302 anti-mouse Hashtag 2 Antibody; Clones M1/42, 30-F11 | BioLegend | Cat# 155803; RRID: AB_2750033 |
| TotalSeq™-A0307 anti-mouse Hashtag 7 Antibody; Clones M1/42, 30-F11 | BioLegend | Cat# 155813; RRID: AB_2750039 |
| TotalSeq™-A0308 anti-mouse Hashtag 8 Antibody; Clones M1/42, 30-F11 | BioLegend | Cat# 155815; RRID: AB_2750040 |
| TotalSeq™-A0309 anti-mouse Hashtag 9 Antibody; Clones M1/42, 30-F11 | BioLegend | Cat# 155817; RRID: AB_2750042 |
| TotalSeq™-A0310 anti-mouse Hashtag 10 Antibody; Clones M1/42, 30-F11 | BioLegend | Cat# 155819; RRID: AB_2750043 |
| TotalSeq™-A0311 anti-mouse Hashtag 11 Antibody; Clones M1/42, 30-F11 | BioLegend | Cat# 155821; RRID: AB_2750136 |
| TotalSeq™-A0312 anti-mouse Hashtag 12 Antibody; Clones M1/42, 30-F11 | BioLegend | Cat# 155823; RRID: AB_2750137 |
| **Biological samples** | | |
| Peripheral blood mononuclear cells (PBMCs) from indicated individuals | This manuscript | N/A |
| **Chemicals, peptides, and recombinant proteins** | | |
| RPMI 1640 Medium | Life Technologies | Cat# 11875-119 |
| Foetal Bovine Serum, NZ origin, heat inactivated | Assay Matrix | Cat# ASFBS-HI-NZ |
| Ionomycin calcium salt | Sigma Aldrich | Cat# I0634-1MG |
| **Critical commercial assays** | | |
| Foxp3/Transcription Factor Staining Buffer Set | Thermo Fisher | Cat# 00-5523-00 |
| NovaSeq S4 230 cycles kit | Illumina | Cat# 20447086 |
| AllPrep DNA/RNA Mini Kit | Qiagen | Cat# 80204 |
| Chromium Next GEM Single Cell 3’ Kit v3.1, 16 rxns | 10X Genomics | Cat# PN-1000268 |
| **Deposited data** | | |
| Raw sequence data, from single-cell RNA sequencing of sorted mouse B cell populations | This manuscript | ENA PRJEB49382 |
| **Experimental models: Organisms/strains** | | |
| Mouse: *Stat3^T716M^:* C57BL/6-Stat3<em1>/Ausb | This manuscript - generated by Prof. Robert Brink | N/A |
| Mouse: *Stat3^K658N^:* C57BL/6-Stat3<em2>/Ausb | This manuscript - generated by Prof. Robert Brink | N/A |
| Mouse: C57BL/6J: C57BL/6 JAusb | The Jackson Laboratory | JAX: 000664 |
| Mouse: CD45.1: B6.JSL-*Ptprc^a^Pepc^b^*/BoyJAusb | The Jackson Laboratory | JAX: 002014 |
| Mouse: *Rag1^KO/KO^*: B6.129S7-*Rag1^tm1Mom^*/JAusb | The Jackson Laboratory | JAX: 002096 |
| **Software and algorithms** | | |
| GraphPad Prims 9 – Version 9.0.0 | GraphPad | <https://www.graphpad.com> |
| FlowJo V10 – Version 10.4.0 | FlowJo | <https://www.flowjo.com> |
| *Limma* | (Ritchie et al., 2015) | <https://bioconductor.org/packages/release/bioc/html/limma.html> |
| *SAVER* | (Huang et al., 2018) | <https://github.com/mohuangx/SAVER> |
| CellRanger | 10X Genomics | Cell Ranger Single Cell v.2.0 |
| Seurat | (Satija et al., 2015) | <https://github.com/satijalab/seurat/> |

RITCHIE, M. E., PHIPSON, B., WU, D., HU, Y., LAW, C. W., SHI, W. & SMYTH, G. K. 2015. limma powers differential expression analyses for RNA-sequencing and microarray studies. *Nucleic Acids Res,* 43**,** e47.

SATIJA, R., FARRELL, J. A., GENNERT, D., SCHIER, A. F. & REGEV, A. 2015. Spatial reconstruction of single-cell gene expression data. *Nat Biotechnol,* 33**,** 495-502.
